# Tumour microenvironments impose translational repression that limits natural killer cell persistence

**DOI:** 10.64898/2026.07.06.736719

**Authors:** Cristhiane Favero Aguiar, Maxim A. Nosenko, Carrie Corkish, Cathal Keane, Yuliya Skabytska, Clair M. Gardiner, Lorraine Brennan, Linda V. Sinclair, David K. Finlay

**Author notes:** **Corresponding Author:** David Finlay, School of Biochemistry and Immunology, Trinity Biomedical Sciences Institute, Trinity College Dublin, 152-160 Pearse Street, D02R590, Dublin, Ireland. These authors contributed equally to this work (Cristhiane Favero Aguiar, Maxim A. Nosenko). Funding received from: - Research Ireland - A Enterprise Ireland innovative partnership with ONK Therapeutics - Marie Skłodowska-Curie Actions.

## Abstract

Natural killer (NK) cells infiltrate many solid tumours, yet the mechanisms that determine their functional heterogeneity across tumour types remain poorly understood. Differences in tumour immunogenicity, inhibitory signalling, and nutrient availability have all been implicated, but unifying explanations are lacking. Here, we compared four syngeneic tumour models implanted at identical anatomical sites to isolate tumour-intrinsic effects on NK-cell fate. Tumour-infiltrating NK cells displayed striking tumour-specific differences in cytokine production, cytotoxic protein expression, and persistence. These differences were not explained by cytokine availability or global features of the tumour metabolic environment. Instead, quantitative proteomics and time-resolved *in vivo* labelling revealed that NK cells enter tumours in a functionally competent state but rapidly diverge thereafter. In suppressive tumour microenvironments, NK cells undergo early mitochondrial loss, translational repression, and impaired proteostatic responses, accompanied by increased apoptotic priming. These defects result in reduced effector function and failure of intratumoural persistence despite preserved recruitment. In contrast, permissive tumours sustain NK-cell translational capacity, cytokine responsiveness, and long-term residency. Together, these findings identify disruption of translational and mitochondrial homeostasis as a central mechanism limiting NK-cell persistence in solid tumours. This work establishes early tumour-induced defects in protein synthesis and cellular fitness as key constraints on durable NK-cell immunity and provides a framework for restoring effective innate anti-tumour responses.

## Introduction

Natural killer (NK) cells are innate lymphocytes that recognise infected and transformed cells for killing using germline-encoded activating and inhibitory receptors^1,2^. NK-cell effector function is shaped by cytokines, influenced further by stress ligands, and the local metabolic environment, and is frequently impaired in solid tumours^3-5^. This dysfunction can manifest as reduced cytotoxicity, diminished cytokine production, and altered receptor expression profiles. However, the mechanisms by which tumour microenvironments (TMEs) impose dysfunction on NK cells remain incompletely resolved, particularly at the level of coordinated metabolic regulation.

The TME can constrain NK-cell metabolism through hypoxia, nutrient limitation, and tumour-derived metabolites^6,7^. Amino acid transport, mTORC1 signalling, and mitochondrial function are central to NK-cell activation and cytotoxic granule biogenesis, and are sensitive to extrinsic cues such as TGF-β and adenosine^8-14^. Beyond substrate availability, metabolic stress can also impair NK-cell function by disrupting proteostasis, organelle integrity, and cellular stress-response pathways^15-17^. While these mechanisms are increasingly appreciated, studies have typically focused on one or a few tumour models, and it remains unclear whether distinct TMEs impose shared or tumour-specific metabolic constraints on NK cells.

In this study, we directly address this question using a controlled comparative framework. We examined four syngeneic tumour models (B16, MC38, LLC, and E0771) implanted subcutaneously at the same anatomical site and profiled tumour-infiltrating NK (TiNK) cells using quantitative proteomics, tumour-interstitial metabolomics, functional assays, and time-resolved *in vivo* labelling. This approach allowed us to define how distinct TMEs shape NK-cell metabolic state, proteostatic capacity, and persistence.

We find that NK cells adopt fundamentally distinct states across tumour types. While MC38 and E0771 tumours support metabolically resilient and functionally competent NK populations, B16 and LLC TMEs induce early mitochondrial loss, marked translational repression, and impaired stress-response engagement. These changes are associated with reduced cytokine production and cytotoxic capacity and occur despite preserved cytokine availability and intact NK-cell recruitment. Instead, NK-cell dysfunction in these settings is linked to disruption of translational and mitochondrial homeostasis following tumour entry.

## STAR Methods

### Mice

Male and female C57BL/6JOlaHsd mice (8–12 weeks, 19–25 g; Inotiv) were housed under specific pathogen-free conditions in individually ventilated cages (≤5 mice per cage) with environmental enrichment, ad libitum chow and water, and a 12-h light/dark cycle under institutional temperature and humidity standard operating procedures. Mice were acclimatised for ≥1 week and monitored daily (twice daily near endpoints). Softened food and hydrogel were provided as needed. All procedures were approved by the Health Products Regulatory Authority and the Trinity College Dublin Animal Research Ethics Committee (AE19136/P157).

### In vivo tumour models and treatments

The experimental unit was one mouse unless otherwise stated. For proteomics, each biological replicate consisted of pooled tumour-derived material from 3–8 mice. In cases of low cell recovery, additional pooling of sorted fractions derived from the same tumour group was performed after cell sorting to obtain sufficient material for analysis (Supplementary Table 1). Tumours (B16-F10, LLC, MC38, EO771) were implanted subcutaneously and analysed at day 14 unless indicated otherwise. Additional experiments included tumour time-stamping (anti-CD45 labelling), adoptive transfer of NK cells and poly(I:C) stimulation. Sample sizes were based on prior experience. Exclusion criteria included tumour ulceration, illness, failed tumour take, early euthanasia, or flow cytometry QC failure (<50% viable TILs or <100 NK cells). Mice were randomised; acquisition and processing were balanced across groups. Blinding was not feasible, but gating used predefined templates and FMO controls.

Time-in-TME labelling was performed using sequential intravascular anti-CD45 staining. Mice received intravenous injections of fluorophore-conjugated anti-CD45 antibodies at defined timepoints (e.g. 72 h, 48 h, and 24 h prior to tissue harvest), allowing temporal labelling of circulating NK cells entering tumours. At endpoint, tumour-infiltrating NK cells were identified based on CD45 label combinations corresponding to defined residence times within the tumour microenvironment. Gating strategies were standardised across experiments using appropriate flow cytometry controls (Supplementary Fig. 1).

### Tumour cell lines and implantation

MC38, EO771, LLC, B16-F10 (ATCC) were cultured in DMEM + 10% FBS, 1% penicillin/streptomycin, 2 mM glutamine. Tumours were established by s.c. injection of 0.5x10^6^ cells in 100 µL PBS. Tumours were measured every 2 days; volume was calculated as V = (L x W^2^)/2. Mice were euthanised at day 14 or upon reaching humane endpoint criteria.

### Poly(I:C) stimulation

Mice received 200 µg poly(I:C) (InvivoGen) (i.p.) or PBS control. Tissues were collected 24 h later.

### Flow cytometry

Tumours were dissociated using the Miltenyi dissociation kit and GentleMACS, followed by leukocyte enrichment (70/40% Percoll). Spleens were mechanically dissociated and RBC-lysed. Cells were Fc-blocked and stained (1:200; Supplementary Table 2) for 25 min (4 °C). Dead cells were excluded using Zombie NIR. Data were acquired on BD LSRFortessa or Agilent Novocyte instruments and analysed in FlowJo. Mature NK cells were gated as live CD45⁺NK1.1⁺dump⁻ (Supplementary Fig. 1A). MFI values were normalised to naïve splenic NK cells within each batch.

### Functional and metabolic assays

Mitochondrial mass and membrane potential were assessed using nonyl acridine orange (100 nM) or MitoTracker Red CMXRos (200 nM) incubated for 20 min at 37 °C prior to surface staining. For *ex vivo* stimulation, tumour, blood, or spleen suspensions were cultured in RPMI supplemented with 10% fetal bovine serum and stimulated with PMA (100 ng/mL) and ionomycin (500 ng/mL) in the presence of Brefeldin A and Monensin for 3 h. Where indicated, 2-deoxyglucose (50 mM) or oligomycin (1 µM) was added 15 min before stimulation. Cells were fixed, permeabilised, and stained for intracellular IFN-γ, granzyme B, and perforin. Slc1a5-mediated uptake was measured using L-homopropargylglycine^18^ (HpG, 400 µM, 5 min), and nascent protein synthesis was determined using O-propargyl-puromycin (OPP, 20 µM, 20 min) incorporation, followed by fixation, permeabilisation, and click-chemistry detection with Alexa Fluor 488 azide.

### Cell sorting

Cells were stained as described and sorted on a BD FACSAria Fusion. NK cells were sorted directly into HBSS, centrifuged, washed, pelleted and snap-frozen. Some samples were pooled to obtain sufficient cell numbers for downstream assays.

### Metabolomics and cytokine quantification

Tumour interstitial fluid was collected by centrifugation through nylon mesh; serum was obtained by cardiac puncture and centrifugation (Supplementary Table 3). Targeted metabolomics was performed using the Biocrates Q500 platform, with LC-MS/MS and FIA-MS/MS acquisition on a Sciex QTRAP 6500+. Cytokines in tumour interstitial fluid were measured using a customised LegendPLEX bead array (BioLegend).

### Adoptive transfer of NK cells into B16-F10 tumour bearing mice

Splenic NK cells were expanded for 7 days in IL15 (10ng/ml) as previously described ^9^. NK cells were purified and activated with IL18 alone or IL18 (50ng/ml) plus IFNβ (100U/ml) for 18 h. 2x10^6^ NK cells in 50µl were injected s.c. at the base of B16-F10 tumours on day 3, 7 and 10 after tumour injection. Tumours were harvested on day 15 and tumour volume and tumour NK cells quantified.

### Proteomic analysis of tumour-infiltrating NK cells

#### Sample preparation and mass spectrometry

Snap-frozen pellets of NK-cells, sorted from tumours, or spleens of mice 24h after i.p. injection with PBS or Poly(I:C) (Supplementary Table 1), were lysed in 5% SDS, 50 mM triethylammonium bicarbonate (TEAB), and protein concentration was determined by micro-BCA assay. Aliquots containing 100 µg protein were processed using the S-Trap mini protocol (Protifi), including reduction with dithiothreitol, alkylation with iodoacetamide, and two-step tryptic digestion (1:40 enzyme-to-protein ratio, 37 °C). Peptides were sequentially eluted, dried, and stored at -20 °C prior to analysis.

Approximately 1.2 µg peptide per sample was analysed by nano-LC on an UltiMate 3000 RSLCnano system coupled to an Orbitrap Exploris 480 mass spectrometer (Thermo Fisher). Peptides were separated on a 75 µm x 50 cm C18 analytical column using a 120 min 3-35% acetonitrile gradient at 300 nL min^-1^ and ionised by an EASY-Spray source in positive mode. Data were acquired in data-independent acquisition (DIA) mode with full MS scans over m/z 350-1650 followed by stepped-collision-energy MS/MS scans (Orbitrap resolution 30,000). All data were acquired in profile mode.

#### Protein quantification and preprocessing

Raw data were processed in Spectronaut 14 (directDIA) against the UniProt mouse database. Absolute protein copy numbers were estimated using the proteomic ruler method. Downstream analyses were restricted to proteins with mean copy number ≥50. Proteomic analyses focused on tumour-infiltrating NK (TiNK) cells from MC38, LLC, EO771, and B16 tumours (n = 16). Activated and naïve splenic NK cells were included as a reference for baseline states for certain biological analyses. Unless otherwise stated, protein and metabolite abundance values were log-transformed and z-scored prior to principal component analysis. PCA was performed using prcomp in R with centered and scaled input matrices.

#### Proteome analyses

##### Principle component analysis

Unless otherwise stated, protein and metabolite abundance values were log-transformed and z-scored prior to principal component analysis. PCA was performed using prcomp in R with centered and scaled input matrices. Within-group proteomic variability was quantified as the Euclidean distance of each sample to the centroid of its tumour group in multivariate protein space.

##### Pathway analysis

Differential protein expression was analysed using limma, and results from four TiNK conditions relative to activated NK cells were used for downstream pathway analysis. For each comparison, proteins were ranked by limma t-statistics to generate ordered lists without applying a significance threshold. Gene set enrichment analysis (GSEA) was performed using the fgsea R package with Reactome pathways obtained via msigdbr. Pathways with a false discovery rate (FDR) < 0.05 were considered significantly enriched (Supplementary File Tab 13). The union of significant pathways across all TiNK conditions was used to define a common pathway set. To reduce redundancy, pathways were grouped based on gene overlap using the Jaccard similarity index, followed by hierarchical clustering (average linkage). Clusters were defined by cutting the dendrogram into k = 10 groups and annotated based on pathway composition. Median normalized enrichment scores (NES) across pathways were used to summarise cluster activity per condition. Interferon signalling pathways were analysed separately due to distinct clustering behaviour. NES values for interferon-related pathways were compared between tumour groups (B16/LLC vs E0771/MC38) using a two-tailed unpaired Mann–Whitney test. Interferon signalling activity was quantified using a curated module of canonical interferon-stimulated proteins: Isg15, Ifit1, Ifti3, Oas1a, Oas3, Bst2, Ifi35, Tap1, Tap2, Psmb8, Psmb9 and B2m, calculated as the mean z-scored protein abundance.

##### Translation-centred principal component analysis

A translation-focused principal component analysis (PCA) was performed using curated features representing translational capacity and repression. Feature construction was guided by the functional organisation of the translation machinery. For example, eIF1 abundance was represented by the EIF1 protein alone, as it is an essential limiting factor in complex formation. In contrast, multi-subunit complexes such as eIF2 were modelled considering their required stoichiometry (e.g. the eIF2 trimer), ensuring that feature values reflected functional complex availability rather than individual subunit abundance. Additional features included global translation machinery abundance (eIF, RPL, and RPS proteins), ribosomal subunit balance (RPL vs RPS), and regulatory inhibitory ratios (PDCD4:eIF4A and 4EBP:eIF4E). All features were z-scored prior to PCA. Principal component scores for PC1 and PC2, which together accounted for ∼77% of translational variance, were retained for downstream analyses.

##### Translation deviation score

To quantify tumour-induced translational reprogramming independently of any single PCA axis, a translation deviation score was defined for each tumour sample as the Euclidean distance from the centroid of splenic NK samples in PC1-PC2 space: Translation deviation = sqrt((PC1 - PC1_spleen)^2 + (PC2 - PC2_spleen)^2). This metric captures the magnitude of translational divergence from the normal NK translational state while integrating variance from both principal components.

##### Apoptotic wiring and priming

Apoptotic susceptibility was quantified using predefined protein-based modules capturing intrinsic priming, extrinsic apoptotic sensitivity, mitochondrial commitment, and executioner potential. Module construction was guided by established apoptotic signalling architecture. Intrinsic apoptotic priming was assessed using BIM:BCL-xL (BCL2L11:BCL2L1) and, where applicable, BIM:MCL1 ratios. Mitochondrial commitment was represented by APAF1, AIFM1, and DIABLO, while extrinsic apoptotic sensitivity was captured using TRADD and CASP8. Execution potential was quantified using CASP3 and CASP7, and BID abundance was treated as an independent bridging metric linking intrinsic and extrinsic pathways.

##### Stress modules

Mitochondrial and cellular stress responses were quantified using curated protein modules. Unless otherwise stated, module scores were calculated as the mean z-scored abundance of proteins within each module. Endoplasmic reticulum stress and unfolded protein response (ER-UPR) were assessed using canonical folding and stress-response proteins, including HSPA5, HSP90B1, PDIA3/4/6, ERO1A, HERPUD1,, ATF6, and DDIT3. Oxidative stress responses were quantified using redox and antioxidant proteins, including NQO1, GCLC/GCLM, HMOX1, TXNRD1, SRXN1, GPX1/4, PRDX1–3, and SOD1/2. Hypoxia signatures were defined using glycolytic and hypoxia-responsive proteins, including SLC2A1, HK2, LDHA, PDK1, PGK1, ENO1, ALDOA, VEGFA, CA9, and HIF1A. Amino acid starvation responses were assessed using transport and integrated stress-response components, including SLC1A5, SLC7A5, ASNS, PSAT1, PHGDH, SHMT2, EIF2AK4, ATF4, and PPP1R15A. Proteotoxic stress was quantified using cytosolic chaperones and co-chaperones, including HSP90AA1, HSP90AB1, HSPA1A/B, HSPA8, HSPH1, DNAJB1, and STIP1. Mitochondrial stress pathways were further resolved into functional modules. Mitochondrial unfolded protein response (UPRmt) and matrix proteostasis were assessed using HSPD1, HSPE1, CLPP, CLPX, LONP1, HSPA9, and HTRA2. Antioxidant and redox control was quantified using PRDX3, SOD2, and TXN2. Inner mitochondrial membrane protease activity was represented by YME1L1, OMA1, PMPCA, and PMPCB, while cytosolic co-chaperone activity was quantified using HSPE1, DNAJA3, and HSPH1. Mitochondrial assembly and metabolic support pathways included IDH2, COX17, NDUFAF1, ATP5IF1, SHMT2, and auxiliary complex I/IV assembly factors where detected.

### Statistical analysis

All analyses were performed in R (v4.5.3) and GraphPad Prism (v9). Differential expression: limma; Correlations: Spearman and Pearson; Multivariable models: linear regression ; PCA: dimensional reduction; Multiplicity was controlled by Benjamini–Hochberg FDR. Normality and homogeneity of variance were assessed using Shapiro-Wilk and Levene/Brown–Forsythe tests. Cell-based assays used ANOVA or non-parametric equivalents as appropriate. Outlier handling was pre-specified and applied consistently across datasets. Significance was defined as p < 0.05.

## Results

### NK cell dysfunction is prominently observed in B16 tumours

To assess how distinct tumour microenvironments shape NK cell responses, we compared four syngeneic tumour models representing immunogenic (MC38, E0771) and less immunogenic (B16, LLC) settings. All tumours formed palpable masses over ∼15 days (Fig. 1A). Tumour immune composition varied markedly between models (Fig. 1B, C). While macrophages dominated the infiltrate across all tumours, B16 tumours showed reduced immune-cell density and were enriched for B cells (Fig. 1D). NK-cell frequencies were highest in MC38 tumours and significantly lower in B16 and LLC (Fig. 1D).

**Figure 1.**
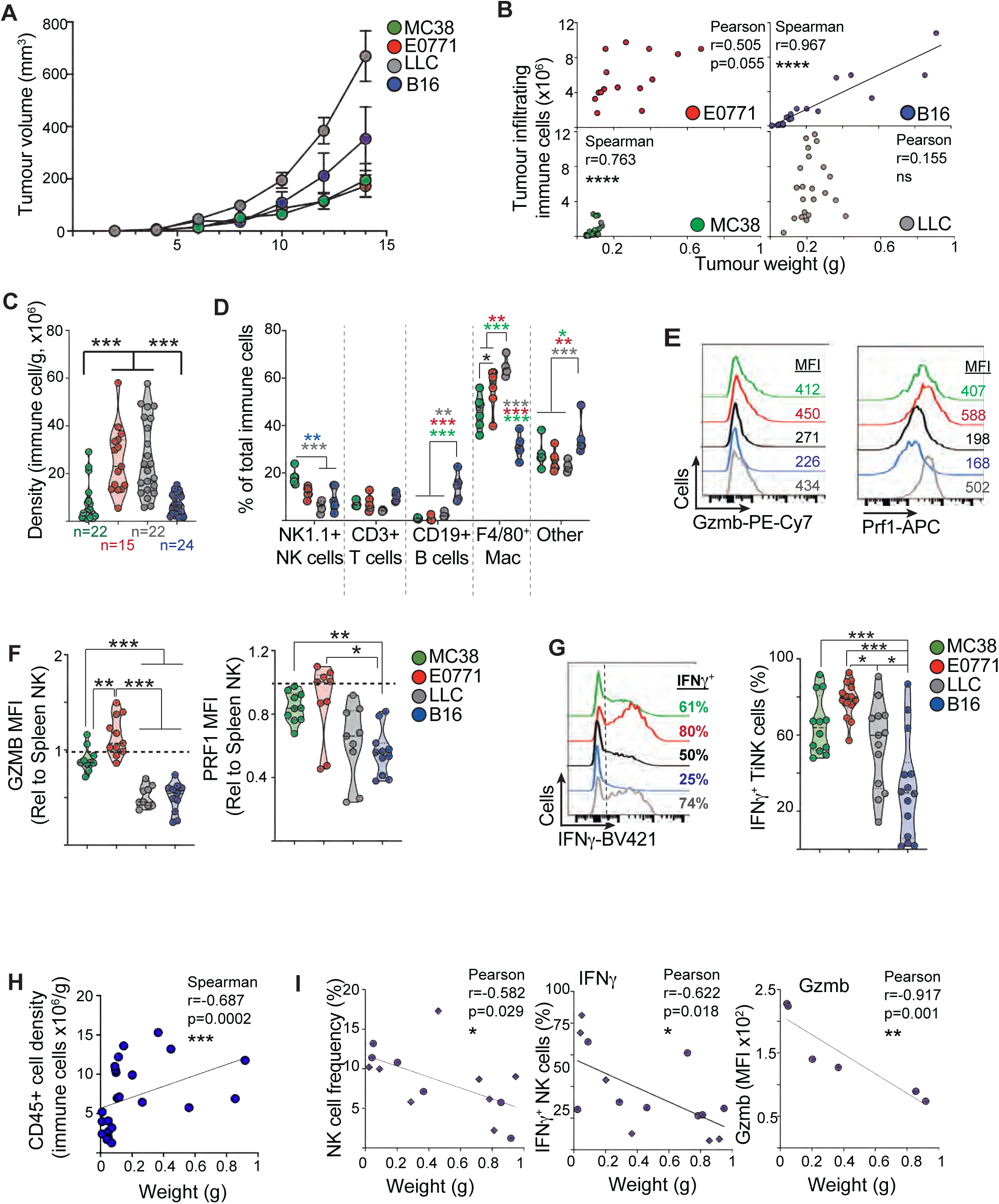
NK cell dysfunction in B16 tumours. (**A-I**) Mice were challenged with MC38 (green), B16 (blue), LLC (grey), or E0771 (red) tumours (s.c. 14-15 days). Tumours were measured and analysed for immune infiltration and immune phenotype by flow cytometry. Splenic NK cells from pooled mice were used as controls.(**A**) Representative tumour growth curves. (**B-C**) Tumour weight, total immune infiltration, and immune density (cells per g tumour; n=15-24). (**D**) Frequency of indicated immune populations among CD45+ tumour cells (n=4-6). (**E-F**) Granzyme B and perforin expression in TiNK and splenic NK cells (n=9-11). (**G**) IFNγ expression in TiNK versus splenic NK cells. (H-I) Correlation of immune parameters (total immune density, NK density, granzyme B, IFNγ) with tumour weight in B16 tumours. Data are representative (A, E, G,), or show individual tumours with correlations (B, H, I) or group medians (C, D, F, G,). Data represent individual mice (n = 15–24 [B-C, H-I], 4-14 [E-G] per group), from ≥10 independent experiments. Statistical analysis was performed using ANOVA or non-parametric tests as appropriate, with Tukey (F, G) or Dunn’s (C, D) post-tests. ns, not significant; *p < 0.05; **p < 0.01; ***p < 0.001; ****p < 0.0001.

We next assessed TiNK cell function directly *ex vivo*. Cytotoxic effector expression was reduced in B16 and LLC tumours, with decreased granzyme B in both models and reduced perforin in B16 (Fig. 1E,F). Cytokine production was also impaired, with fewer IFNγ-producing NK cells in B16 compared to all other tumour types and reduced IFNγ in LLC relative to E0771 (Fig. 1G). B16 tumours displayed the greatest variability in both tumour size and NK cell function. Increasing tumour size was associated with increase overall immune cell density but a reduced frequency of NK cells within these immune cells (Fig. 1H,I). Increasing B16 tumour weight was also negatively corrolated to NK cell effector function, including IFNγ production and granzyme B expression (Fig. 1I). Together, these data indicate that B16 tumours impose a pronounced functional impairment on infiltrating NK cells.

### Tumour-specific metabolic states across local and systemic compartments

Cytokine profiling of tumour interstitial fluid (TIF) revealed minimal differences across tumour types, with the exception of increased IFNγ in MC38 tumours (Fig. 2A). We therefore examined whether tumour-specific metabolic environments might distinguish these models.

**Figure 2.**
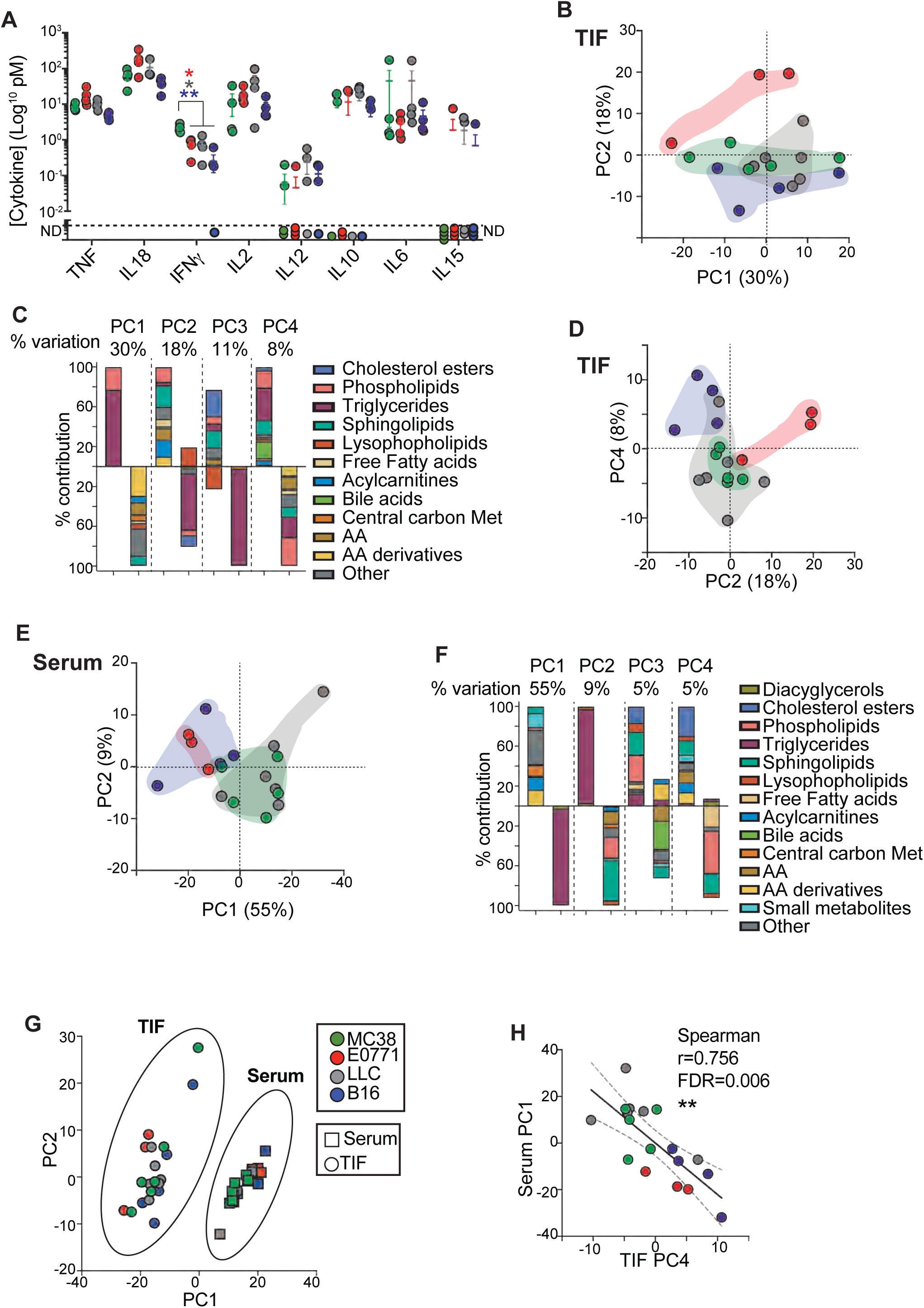
Metabolomic features link the TME to systems metabolism. **(A)** Concentrations of cytokines detected in the tumour interstitial fluids and analysed using cytometry bead array. Dashed line indicates limit of detection. Data are presented as mean±SD, n=4 mice. (**B-I**) Mice were challenged with MC38 (green), B16 (blue), LLC (grey), or E0771 (red) tumours (s.c. 14-15 days). TIF and serum was collected and metabolites quantified. (**B,D,E**) PCA of TIF (B,D) and serum (E) metabolites showing PC1 and PC2 (B,E) and PC2 and PC4 (D). Positive and negative loading metabolites were consolidated to theme that labels each axis. (**C,F**) Categories of metabolites with contributions to TIF(C) and serum (F) PCA PC1 to PC4. % variability captured by each PC is indicated. (**G**) PCA of TIF and serum data together showing PC1 and PC2. (H) Corelation analysis of serum PC1 with TIF PC4). Data is mean +/- SEM (A), individual tumour metabolomic replicates (B,D,E,G,H), mean % contributions to PC (C,F). Data represent individual mice (n = 4 [A], 3-6 [B-H] per group) collected from >2 independent experiments. Statistical analysis was performed using ANOVA with Tukey post-tests. *p < 0.05; **p < 0.01;

Metabolomic profiling of TIF revealed distinct tumour-specific signatures. Glycolytic metabolites were broadly similar across tumours, with uniformly high lactate levels, suggesting that glycolysis does not account for inter-tumour differences (Supplementary File Tab 1). PCA showed that variation along PC1 was driven by lipid versus amino acid-enriched environments, while higher-order components (PC2–PC4) further separated tumour types (Fig. 2B-D, Supplementary File Tab 2).

Individual tumours occupied distinct metabolic regions within PCA space. E0771 tumours were associated with amino acid- and acylcarnitine-rich profiles, whereas B16 tumours displayed a metabolically distinct state characterised by enrichment in lipid-derived signalling molecules and metabolic intermediates. In contrast, MC38 and LLC tumours showed more intermediate metabolic profiles.

To determine whether these tumour-associated metabolic states were reflected systemically, we profiled matched serum samples. Serum PCA revealed separation of tumour types along a primary axis distinguishing triglyceride-rich and metabolite-rich states, with B16 and E0771 exhibiting triglyceride-enriched profiles (Fig. 2E,F). Although TIF and serum profiles were distinct, a significant correlation between TIF PC4 and serum PC1 indicated coupling between local and systemic metabolic states (Fig. 2G,H).

Together, these data show that tumour types are characterised by distinct yet interconnected metabolic environments, without a single shared metabolic feature that explains differences in NK cell function.

### Tumour-infiltrating NK cells acquire distinct proteomic programmes

To define how tumour microenvironments shape NK cell states, we performed quantitative proteomics on TiNK cells from four tumour models alongside naïve and in vivo-activated splenic NK cells (Supplementary Table 1) ^19^. Principal component analysis (PCA) revealed clear separation between naïve and in vivo-activated NK cells, reflecting extensive activation-induced proteome remodelling (Fig. 3A-C). Differential expression analysis of 4565 unique proteins identified 749 upregulated and 97 downregulated proteins following *in vivo* activation (Fig. 3D, Supplementary File Tab 4).

**Figure 3.**
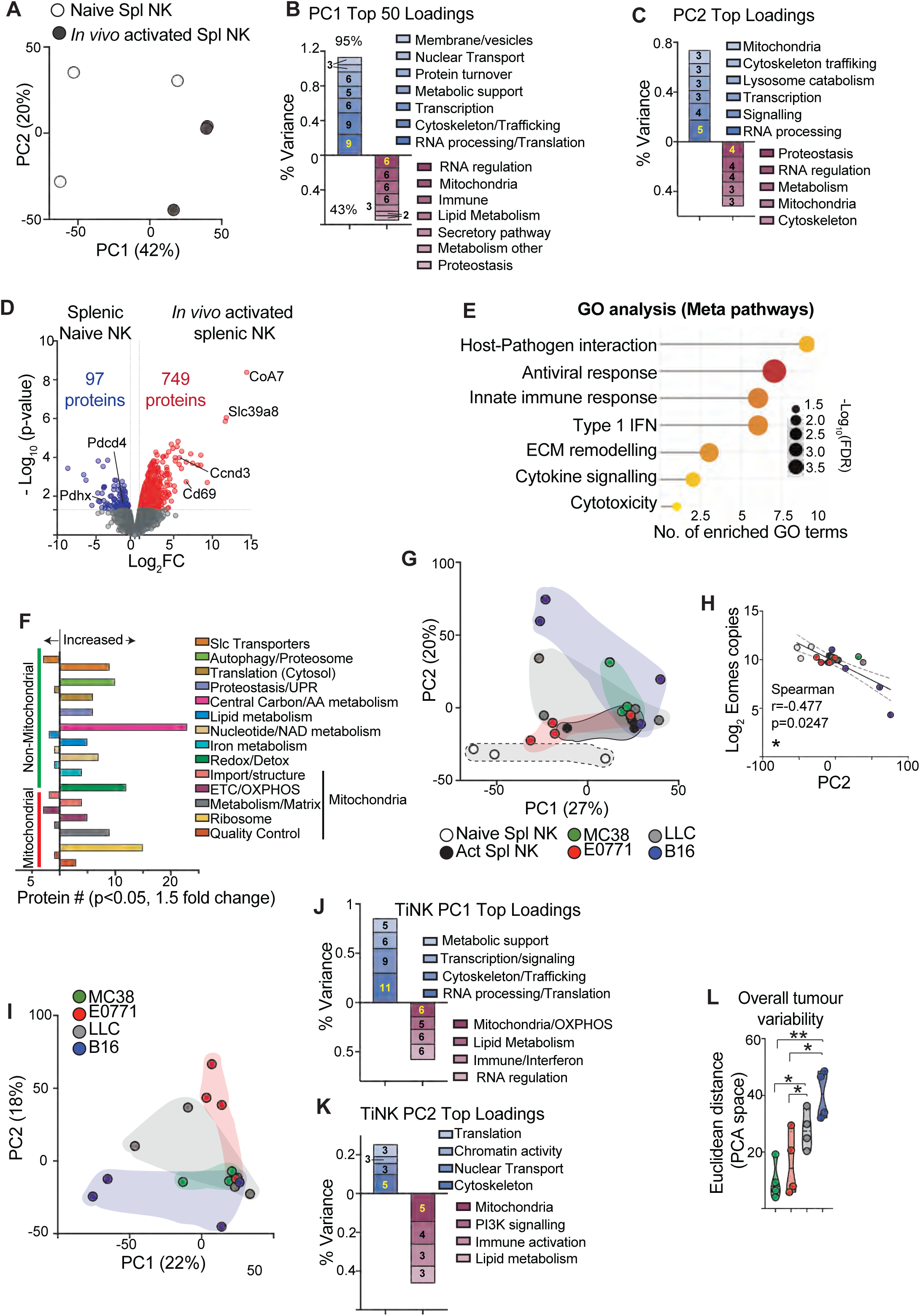
Significant variation is a dominant feature of TiNK cell proteomes. **(A-F)** NK cells were isolated from mice following i.p injection of PBS or Poly(I:C) (200 µg/mouse, *in vivo* activated) and proteome quantified. (**A**) PCA and (**B,C**) categorisation of the top 50 positive and negative loading proteins for PC1 (B) and PC2 (C). (**D**) Volcano plot showing differentially expressed proteins (Limma linear modelling, p<0.05, 1.5 fold change) in naïve NK cells (blue) and *in vivo* activated NK cells (red). **(E)** GO pathway analysis of DE proteins between naïve and *in vivo* activated NK cells. Significantly enriched pathways (FDR<0.05) were manually grouped by related keywords and these meta groupings are shown with significance shown (dot size) and number of related pathways per group (x-axis). (**F**) Mitochondrial and non-mitochondrial metabolic proteins were collated, categorised and numbers of increased or decreased protein for *in vivo* activated NK cells shown. (**G-L**) Alternatively, mice were challenged with MC38 (green), B16 (blue), LLC (grey), or E0771 (red) tumours (s.c. 14-15 days), and TiNK proteomes quantified. (**G**) PCA of naïve, *in vivo* activated splenic NK cells plus TiNKs from each tumour type. (**H**) Correlation of PC2 with Eomes protein copy numbers. (**I**) PCA of TiNK samples only and (**J,K**) categorisation of the top 50 positive and negative loading proteins for PC1 (J) and (L,K). (**L**) Quantification of the Euclidean distance within the PCA space for MC37, E0771, LLC and B16 TiNK. Data is proteomes from 3 (splenic NK) and 4 (TiNK) replicates. Data represent proteomic samples (n = 3 [splenic NK], 4 [TiNK] per group), collected from >2 independent experiments. Data is analysed using PCA (A,G,I), Limma linear modelling and p<0.05 (D,F), GO pathway analysis with FDR <0.05 (E), Spearman correlation (H) and a one way ANOVA with Tukey post-test (O). * p < 0.05; **p < 0.01.

Activated NK cells displayed strong induction of immune and antiviral pathways, alongside marked metabolic remodelling (Fig. 3E,F). This included coordinated upregulation of glycolysis, pentose phosphate pathway, nucleotide biosynthesis, and mitochondrial pathways, consistent with a shift towards an anabolic, biosynthetically active state supporting effector function (Supplementary file Tab 5).

We next compared TiNK proteomes to splenic NK cell states. PCA revealed that TiNK cells occupied a continuum spanning activated and naïve NK cell profiles, with additional separation across PC2 reflecting tumour-specific variation (Fig. 3G). Increased PC2 scores were associated with metabolic and RNA processing pathways and inversely correlated with Eomes protein abundance, suggesting progressive loss of core NK cell identity in subsets of TiNK cells (Fig. 3H, Supplementary File Tab 6).

Analysis of TiNK cells alone revealed substantial inter-tumour heterogeneity, with B16 and LLC samples showing significantly greater variation than MC38 and E0771 (Fig. 3I). Protein loading analysis indicated that variation across principal components was driven by coordinated contributions from RNA processing, mitochondrial function, immune signalling, and lipid metabolism (Fig. 3J-K; Supplementary Fig. 2, Supplementary File Tab 7-10). Consistent with this, Euclidean distance analysis demonstrated increased dispersion of TiNK proteomes in B16 and LLC tumours relative to MC38 and E0771, indicating greater within-tumour variability (Fig. 3I). Together, these data show that tumour microenvironments impose distinct and heterogeneous proteomic states on infiltrating NK cells.

### Reduced type-1 interferon signalling B16 and LLC TiNK cells

To define tumour-specific alterations in NK cell state, we compared TiNK proteomes from each tumour model to in vivo-activated splenic NK cells. Differential expression analysis revealed largely tumour-specific changes, with limited overlap between models (Fig. 4A; Supplementary Fig. 2C,D, Supplementary File Tab 11). Among shared features, the transcription factor Tcf7 was consistently upregulated and inversely correlated with granzyme B abundance across TiNK cells (Fig. 4B).

**Figure 4.**
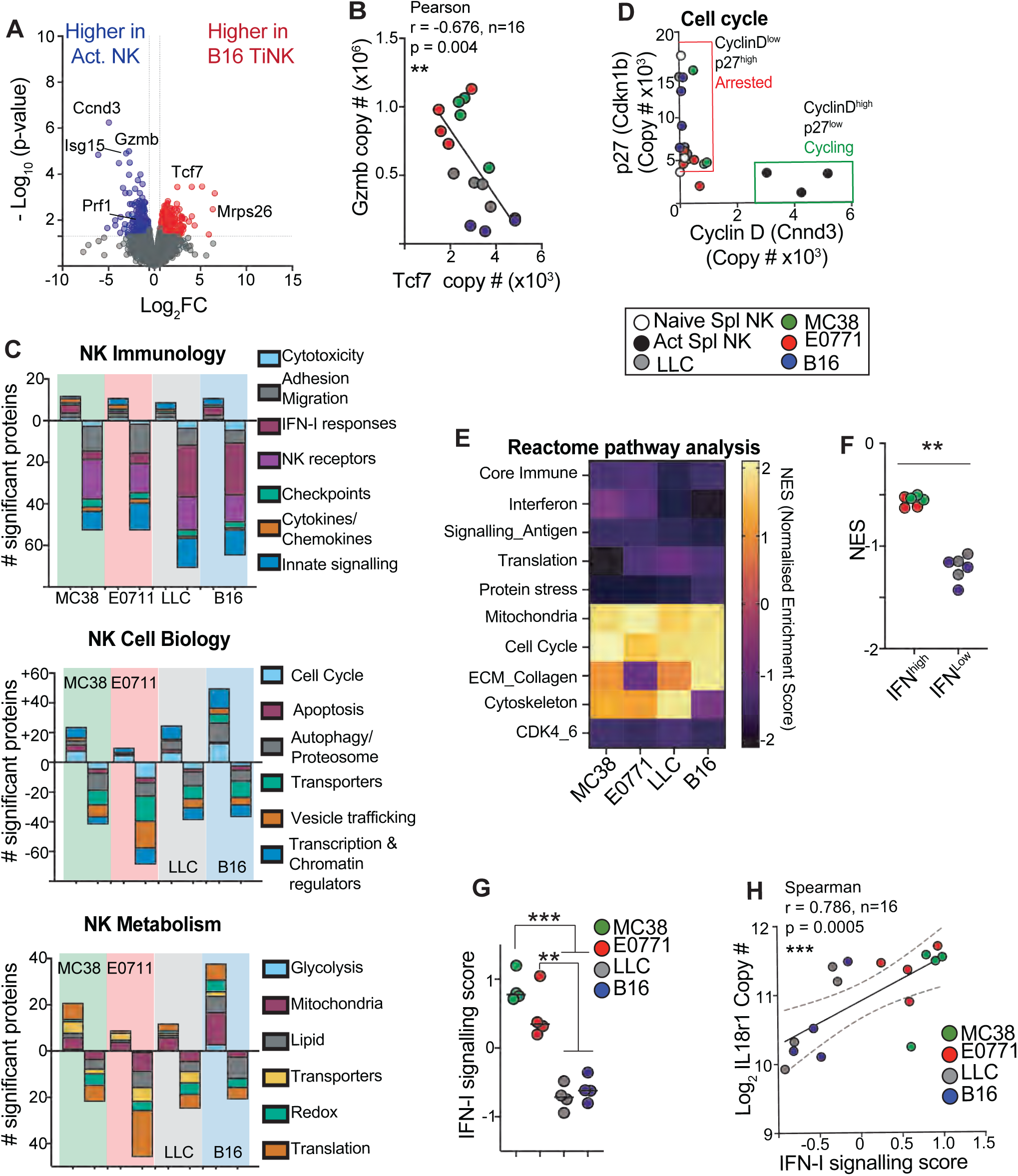
TiNK proteomes identify reduced IFN-I signalling in LLC and B16 tumours. Mice were challenged with MC38 (green), B16 (blue), LLC (grey), or E0771 (red) tumours (s.c. 14-15 days). Alternatively, NK cells were isolated from mice following i.p injection Poly(I:C) (200 µg/mouse, *in vivo* activated). Splenic NK and TiNK proteomes were quantified. **(A)** Example volcano plot showing differentially expressed (DE) proteins (Limma linear modelling, p<0.05, 1.5 fold change) from *in vivo* activated NK cells (blue) or B16 TiNK cells (red). (**B**) Correlation of Gzmb and Tcf7 protein copy numbers for TiNK samples (**C**) DE proteins in pairwise Limma analyses between TiNK and *in vivo* activated NK cells curated into categories relating to (top to bottom panels) NK immunology, NK cell biology and NK metabolism. (**D**) Plot of Cyclin D (x axis) versus p27 (y-axis) protein copy numbers showing distribution of cycling (green box) and arrested (red box) NK cells. **(E)** DE proteins from each pairwise Limma were integrated and analysed by Reatome pathways analysis. Outputs were subjected to hierarchic clustering (k=10) and normalised enrichment scores shown. Orange-yellow; pathways increased in TiNK vs *in vivo* activated NK cells, Purple-black are pathways decreased in TiNK vs *in vivo* activated NK cells). (**F**) NES values for interferon-related pathways (3 separate Reactome outputs) were compared between TiNK groups (B16/LLC vs E0771/MC38). (**G**) Proteomic data for 12 IFN-I regulated proteins combined to compute IFN-I signalling score for each tumour replicate. (**H**) Correlation of IFN-I signalling score with Log2 normalised IL18r1 protein copy numbers. Data represent proteomic samples (n = 3 [splenic NK], 4 [TiNK] per group), collected from >2 independent experiments. Data is analysed using Limma linear modelling and p<0.05 (A) Pearson and Spearman correlations (B, H), Reactome pathway analysis and FDR <0.05 (E,F), a Mann-Whitney unpaired t-test (F), and a one-way ANOVA with Tukey post-test. **, p < 0.01; ***, p < 0.001 NES, Normalised enrichment score.

Functional annotation of differentially expressed proteins revealed broad remodelling of NK cell programmes. Downregulation of immune signalling and cytotoxicity-associated proteins was most pronounced in B16 and LLC TiNK cells, whereas MC38 and E0771 retained higher expression of these pathways (Fig. 4C, Supplementary File Tab 12). In contrast, proteins associated with cell cycle regulation and metabolism showed bidirectional changes across all tumour models, consistent with a shared shift away from the activated NK cell state (Fig. 4D).

Pathway analysis confirmed a global reduction in immune and translational programmes in TiNK cells relative to activated NK cells, alongside increased activity of mitochondrial and cell cycle pathways (Supplementary File Tab 13). While several pathway modules varied between individual tumour types, interferon signalling pathways most closely tracked the functional differences observed across TiNK subsets (Fig. 4E).

Although interferon pathway activity was reduced in all TiNK cells, this reduction was markedly greater in B16 and LLC compared to MC38 and E0771 (Fig. 4G,H). An interferon-response module score further demonstrated substantially lower IFN-I signalling in B16 and LLC TiNK cells. Consistent with this, IFN-I signalling positively correlated with IL-18 receptor abundance, indicating that reduced interferon signalling is associated with diminished cytokine responsiveness in these tumours (Fig. 4H) ^20-22^.

Together, these data identify reduced type I interferon signalling as a defining feature of dysfunctional TiNK cells in B16 and LLC tumours.

### Translational reprogramming in the TME suppresses NK effector programs

Pathway analysis indicated disruption of translational machinery in tumour-infiltrating NK (TiNK) cells. Consistent with this, B16 and LLC TiNK cells showed reduced expression of key translation-initiation components, including eIF1 and stress-responsive kinases, alongside increased expression of translational inhibitory factors (Fig. 5A). In contrast, ribosomal subunit abundance and balance were largely preserved, although increased variability was observed between TiNK samples, particularly in B16 tumours (Fig.5B).

**Figure 5.**
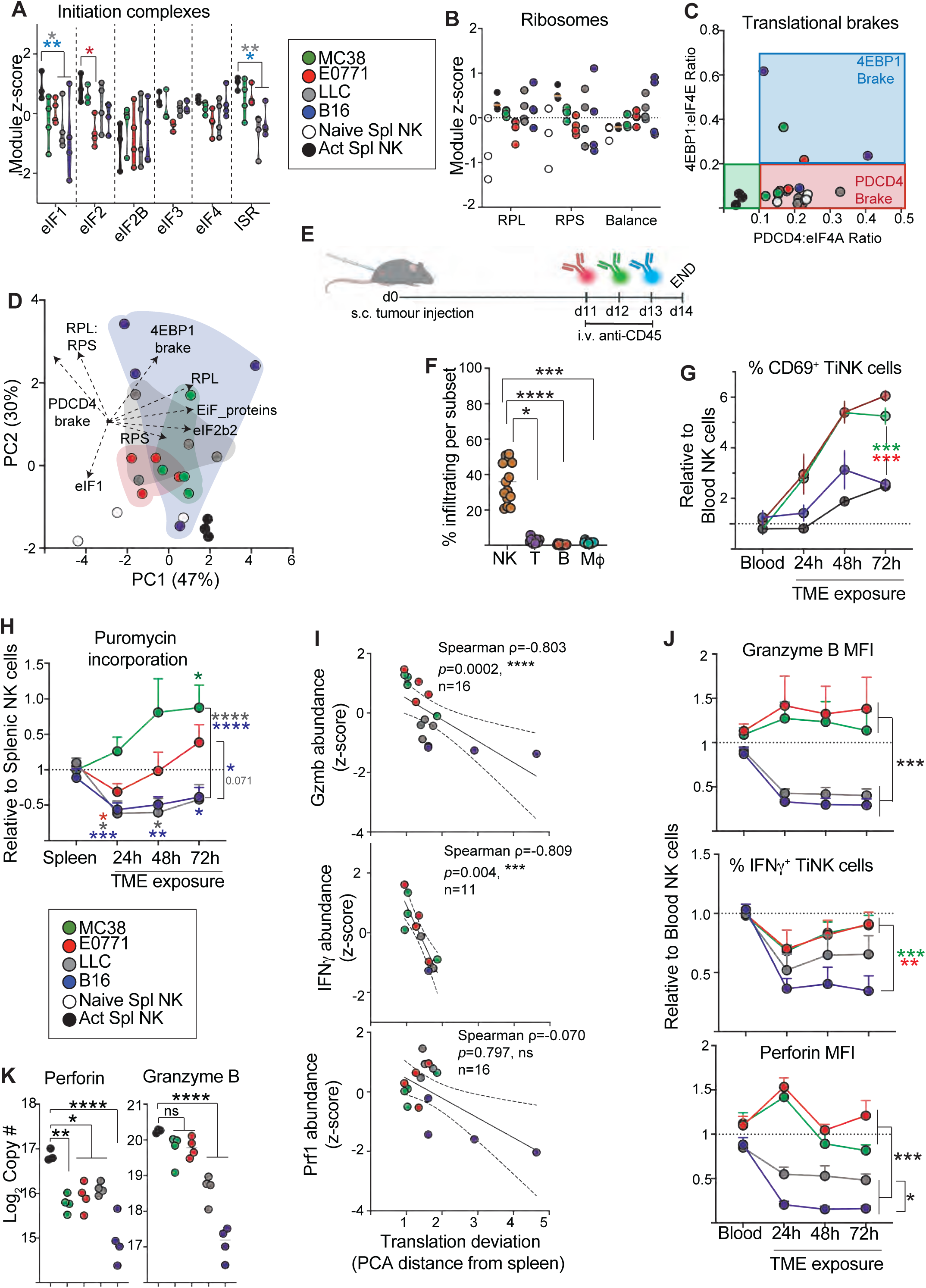
Deficiencies in protein synthesis in B16 and LLC TiNK cells. Mice were challenged with MC38 (green), B16 (blue), LLC (grey), or E0771 (red) tumours (s.c. 14-15 days). Alternatively, NK cells were isolated from spleen of mice following i.p injection PBS (naïve NK) or Poly(I:C) (200 µg/mouse, *in vivo* activated). Splenic NK and TiNK proteomes were quantified. (**A,B**) Module z-scores for translation initiation complexes EIF1-4 and the EIF2 kinases (Perk, Pkr, Gcn2 and Hri) activated by integrated stress response (ISR) (A) and small (RPS) and large ribosome complexes (RPL) plus RPS to RPL balance (B). (**C**) Plot of translational brake mechanism; the EIF4A:PDCD4 (x-axis) and EIF4E:4EBP1 (y-axis) ratios. (**D**) Variation in translational the elements indicated was analysed using a translation focused PCA. (**E-H**) Mice were challenged with MC38 (green), E0771 (red), LLC (grey) and B16 (blue) tumours and on day 11, 12 and 14 received i.v. injection of anti-CD45, each conjugated to a different fluorophore, before isolation and analysis of TiNK on day 15 by flow cytometry. (**E**) Diagram outlining experiment. (**F**) Percentage of newly infiltrated immune cells (NK cell, T cells, B cells and Macrophages) identified as positively stained i.v. anti-CD45. (**G,H**) Surface expression of CD69 (G) and the rate of protein translation (H) for TiNK cells exposed to respective TME for 24, 48 and 72 hours, shown relative to blood and splenic NK cells, respectively. (**I**) Correlation of translation deviation score to Gzmb, IFNγ and perforin abundance (z-scores) (**J**) Expression of Granzyme B and Perforin (MFI) or % IFNγ positive NK cells in TiNK exposed to respective TME for 24-72 h in experiment described in (E). (**K**) Log2 copy number of Perforin and Granzyme B in TiNK cell proteome data. Data represent individual proteomic samples (n = 4 [A-D,I,K] per group), mean +/- SEM of independent mice (n = 13 [F], 8-9 [G,H,J] per group) from at least 2 independent experiments. Statistical analysis was performed using ANOVA or non-parametric tests as appropriate Šidák (A,H) or Tukey post-test. * p < 0.05; **p < 0.01; ***p < 0.001; ****p < 0.0001.

Ratios of translational repressors to their targets revealed increased inhibition of translation initiation in TiNK cells. Both PDCD4 and 4EBP1/2 showed elevated relative abundance compared to their respective binding partners, consistent with enhanced restraint on protein synthesis in multiple tumour models (Fig. 5C).

To integrate these features, we performed PCA of translation-related proteins/modules. Variation along PC1 reflected overall translational capacity, whereas PC2 captured features of translational repression. In vivo-activated splenic NK cells clustered tightly within a high-capacity, low-repression state, whereas TiNK cells showed variable displacement along both axes, indicating heterogeneous translational states (Fig. 5D). This heterogeneity was most pronounced in B16 tumours, where individual samples spanned a wide range of translational capacity and repression states.

To determine whether translational repression reflected acute or sustained exposure to the TME, we measured protein synthesis in TiNK cells over time, using an approach to time-stamp NK cells in the blood and tracks them infiltrating into tumours (Fig. 5E; Supplementary Fig. 1). Across all tumour models, NK cells were the predominant immune subset infiltrating into post day 11 established tumours (Fig. 5F). In this model the expression of CD69 on infiltrating NK cells increased over time on all TiNK cells but to different degrees in each TME (Fig. 5G). Apart from MC38 tumours, translation rates were reduced shortly after tumour entry, as measured by O-propargyl-puromycin (OPP) incorporation (Fig. 5H). While translation recovered over time in E0771 tumours and increased in MC38, it remained suppressed in B16 and LLC TiNK cells. Consequently, the magnitude and persistence of translational suppression were greatest in B16 and LLC tumours.

To determine whether translational changes impact NK cell function, we quantified translational deviation in tumour-infiltrating NK (TiNK) cells relative to activated splenic NK cells. In simple terms, this metric reflects how much each TiNK sample differs from activated NK cells in terms of protein synthesis.

Translational deviation was negatively associated with NK cell effector function. Increased deviation correlated with reduced granzyme B and IFNγ abundance, whereas perforin levels were not associated with translational state (Fig. 5I).

Temporal analysis revealed tumour-specific effects on NK cell function. In B16 and LLC tumours, granzyme B, IFNγ, and perforin were rapidly reduced within 24 h of tumour entry. In contrast, MC38 and E0771 tumours showed minimal or transient effects, with recovery of IFNγ over time (Fig. 5J; Supplementary Fig. 3A,B).

Bulk proteomic analysis of TiNK cells, which includes cells exposed to the tumour microenvironment for extended periods, revealed distinct patterns for cytotoxic proteins.

Perforin abundance was uniformly reduced across all tumour types compared with activated splenic NK cells. In contrast, granzyme B was selectively reduced in B16 and LLC TiNK cells (Fig. 5K).

Together, these data show that tumour microenvironments impose variable but often sustained suppression of protein synthesis in TiNK cells, resulting in translational reprogramming that is associated with impaired NK cell effector function, with the strongest and most sustained defects observed in B16 and LLC tumours.

### Mitochondrial stress and reduced metabolic resilience in B16 TiNK

Given that sustained protein synthesis is required to maintain mitochondrial integrity and function, we next examined whether the translational defects observed in TiNK cells were associated with altered mitochondrial homeostasis. Initial analysis showed that global stress-module scores, including ER stress/UPR and proteostasis pathways, were elevated in activated NK cells relative to naïve NK cells, but were reduced across all TiNK populations (Fig. 6A), indicating a shift away from the activated stress response programme. We next focused on mitochondrial-specific stress pathways. B16 TiNK cells displayed significantly elevated mitochondrial stress signatures, including increased mitochondrial UPR (mtUPR) and mitochondrial reactive oxygen species (mtROS) modules compared to other tumour types (Fig. 6B). This increase was primarily driven by proteins involved in matrix proteostasis, suggesting selective activation of this arm of the mtUPR (Fig. 6C,D). These findings align with the metabolomic features observed in B16 tumours.

**Figure 6.**
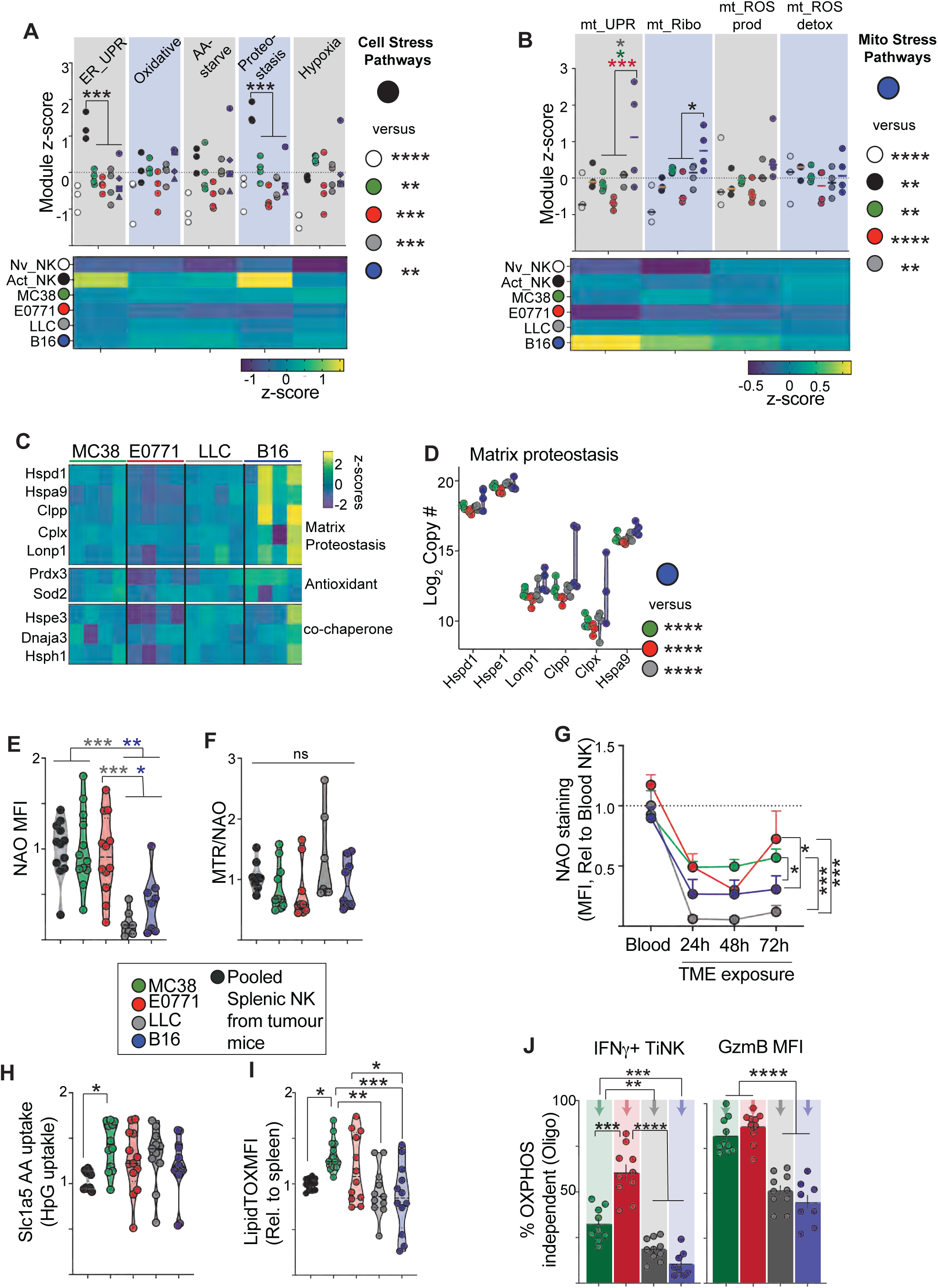
Rapid mitochondrial collapse in tumour infiltrating B16 and LLC TiNK cells. **(A-D)** Mice were challenged with MC38 (green), B16 (blue), LLC (grey), or E0771 (red) tumours (s.c. 14-15 days). Alternatively, NK cells were isolated from spleen of mice following i.p injection PBS (naïve NK) or Poly(I:C) (200 µg/mouse, *in vivo* activated). Splenic NK and TiNK proteomes were quantified. (**A,B**) Module scores for cellular stress pathways (A) and mitochondrial stress pathways (B) shown for each splenic NK and TiNK replicate. The heatmap below summarizes mean module z-scores for each group. **(C, D)** Heatmap showing z-score protein abundance of matrix proteostasis, antioxidant and co-chaperone regulators (C) and log_2_-transformed copy numbers (D) of matrix proteases (Hspd1, Hspe1, Lonp1, Clpp, Clpx and Hspa9) in TiNK cells. (**E-H**) Splenic NK cells or TiNKs were isolated and stained for (E) mitochondrial mass (NAO), (F) mass normalised mitochondrial membrane potential (TMRM/NAO), (H) HpG uptake via AA transporter Slc1a5 and (I) neutral lipid content (LipidTOX). (**G**) Mice were challenged with MC38 (green), E0771 (red), LLC (grey) and B16 (blue) tumours and on day 11, 12 and 14 received i.v. injection of anti-CD45, before isolation and analysis of TiNK on day 15 by flow cytometry. Time dependent changes in mitochondrial mass (NAO), calculated relative to blood NK cells. **(J)** TiNK were isolated from day 14-15 tumours, stimulated for 3 h with PMA/ionomycin +/- oligomycin (1 µM)) and OXPHOS independent IFNγ and granzyme B shown. Data represent mean values and individual proteomic samples (n = 3 [splenic NK] and 4 TiNK] [A-D] per group), individual mice (n= 6-12 [E,F,H,I] per group) and mean +/-SEM of independent mice [n = 5-8 [G], 8-10 [J] per group) from at least 2 independent experiments. Statistical analysis was performed using ANOVA or non-parametric tests as appropriate, with Tukey post-test. * p < 0.05; **p < 0.01; ***p < 0.001; ****p < 0.0001; NAO, 10-N-nonyl acridine orange; HPG, L-Homopropargylglycine; MTR, Mitotracker Red.

Direct assessment of mitochondrial properties revealed reduced mitochondrial mass in B16 and LLC TiNK cells, while membrane potential was largely preserved (Fig. 6E,F). Temporal analysis showed that mitochondrial mass decreased following entry into all tumour microenvironments, but partially recovered in MC38 and E0771, whereas recovery was limited in B16 and LLC TiNK cells (Fig. 6G).

Consistent with these observations, metabolic substrate availability differed across tumour types. Neutral lipid stores were reduced in B16 and LLC TiNK cells compared to other tumours, whereas glutamine uptake capacity showed minimal differences between TiNK populations (Fig. 6H,I).

To assess functional metabolic resilience, TiNK cells were challenged with mitochondrial inhibition. B16 and LLC TiNK cells displayed greater sensitivity to oligomycin, with more pronounced reductions in IFNγ production and granzyme B expression compared to MC38 and E0771 (Fig. 6J). B16 TiNK cells also showed increased sensitivity to glycolytic inhibition, indicating limited metabolic flexibility (Supplementary Fig. 3C).

Together, these data demonstrate that TiNK cells from B16 and LLC tumours exhibit mitochondrial stress, impaired recovery of mitochondrial capacity, and reduced metabolic resilience, consistent with a dysfunctional and metabolically constrained NK cell state.

### Apoptotic priming in B16 and LLC TiNK cells

Given the translational and mitochondrial defects observed in TiNK cells, we next assessed apoptotic priming across tumour models. All TiNK populations showed increased apoptotic priming relative to splenic NK cells, primarily driven by an increased Bim:Bcl_XL_ ratio (Fig. 7A-C). Despite this shared trend, the degree and nature of apoptotic priming varied between tumour types. To resolve this, we analysed intrinsic, extrinsic, and mitochondrial stress–associated apoptotic pathways. This revealed distinct apoptotic profiles across TiNK populations, with B16 TiNK cells exhibiting the greatest heterogeneity, spanning intrinsic, extrinsic, and executional pathways (Fig. 7D). LLC TiNK cells showed a similar but more restricted pattern, whereas MC38 and E0771 TiNK cells displayed more uniform profiles with lower intrinsic priming and reduced evidence of executional activation. These differences indicate that TiNK cells in B16 and LLC tumours are more variably and strongly primed for apoptosis, while those in MC38 and E0771 remain comparatively resistant. This pattern is consistent with the increased metabolic and translational stress observed in B16 and LLC TiNK cells.

**Figure 7.**
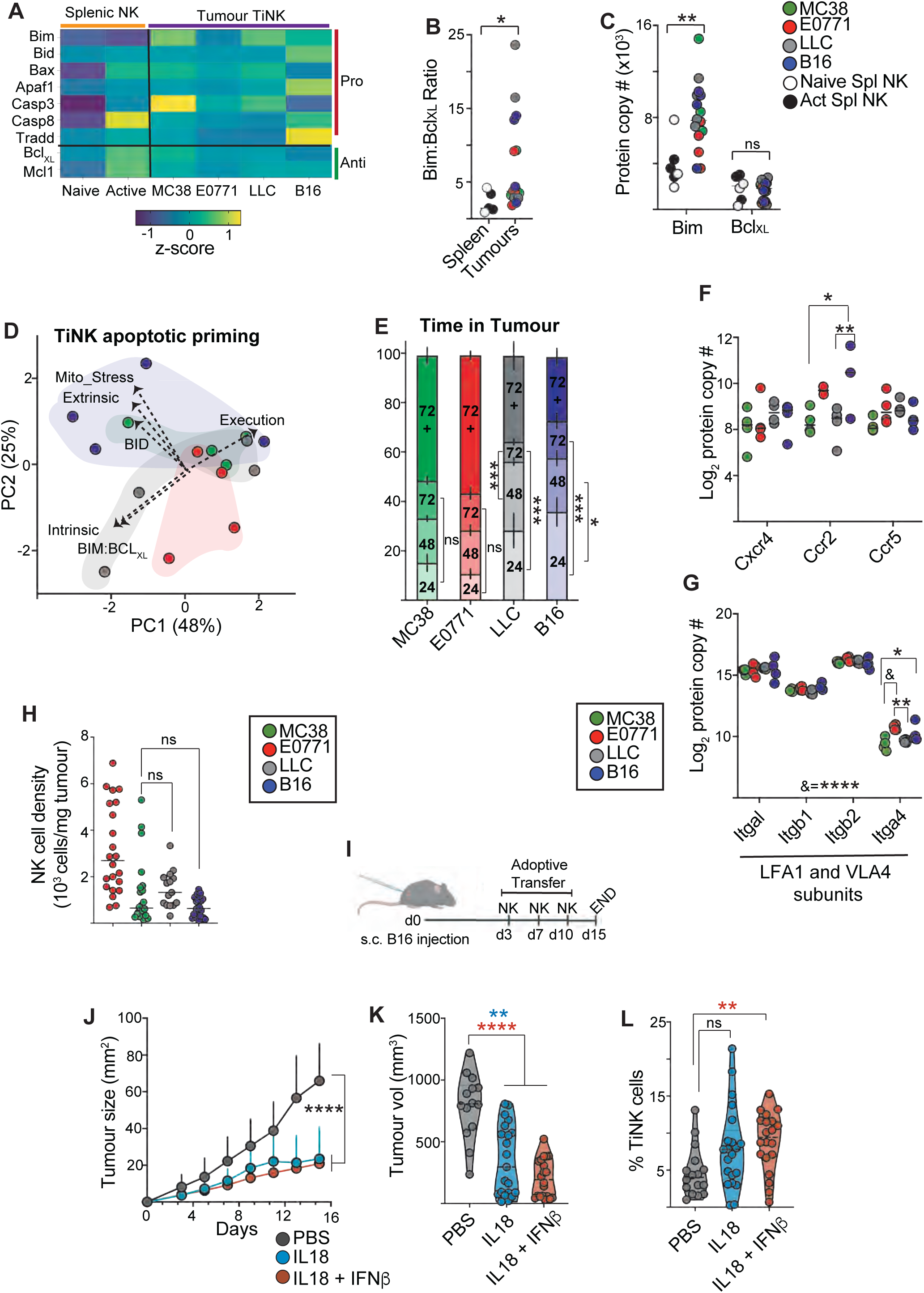
IFN-I/IL-18 link apoptotic priming to TiNK persistence. (**A-C,F,G**) Mice were challenged with MC38 (green), B16 (blue), LLC (grey), or E0771 (red) tumours (s.c. 14-15 days). Alternatively, NK cells were isolated from spleen of mice following i.p injection PBS (naïve NK) or Poly(I:C) (200 µg/mouse, *in vivo* activated). Splenic NK and TiNK proteomes were quantified. (**A,C**) Abundances of apoptosis linked proteins and (**B**) Bim:BclXL ratios. **(D)** Variation in apoptotic priming axis’ in proteome data was analysed using a apoptosis focused PCA. (**E)** Mice were challenged with tumours (s.c. 14-15 days) with i.v. injection of anti-CD45 on day 11, 12 and 13, TiNK were isolated and analysedTiNK on day 14. Time spent within tumours was calculated on the basis of CD45-fluourophore labelling pattern. (**F-G**) Log_2_ normalised protein copy number of chemokine receptors and adhesion molecules on TiNK from each tumour. (**H**) Density of total NK cells per mg of tumour tissue from standard tumour challenge experiments. (**I-L**) Mice were challenged with B16 tumours (s.c) and received 3 intra-tumoral injections of PBS, or *in vitro* expanded NK cells stimulated with IL18 (50 ng/ml) +/- IFNβ (100 U/ml) on days 3, 7, 10 (I). Tumour growth is charted and day 15 tumour volume shown (J,K). Immune cells were isolated and TiNK cell frequency measured by flow cytometry (L). Data represent mean values and individual proteomic samples plus median (n = 3 [splenic NK], 4 [TiNK] [A-C,F-G] per group) collected from a least 2 independent experiments; mean +/- SEM of individual mice (n = 7-9 [E] 16-20 [J] per group), individual mice (n = 15-24 [H] 16-20 [K,L] per group) from 2 [E], 4 [J-L] independent experiments. Statistical analysis was performed using ANOVA or non-parametric tests as appropriate, with Tukey or Šidák (C) post-tests. ns, not significant; * p < 0.05; **p < 0.01; ***p < 0.001; ****p < 0.0001.

Together, these findings identify apoptotic priming as a downstream consequence of tumour-induced metabolic and translational dysfunction, with B16 and LLC tumours driving the strongest pro-apoptotic states in infiltrating NK cells.

### Reduced TiNK persistence in B16 and LLC tumours

The rapid loss of mitochondrial mass observed in B16 and LLC tumours (Fig. 6) coincided with translational repression, mitochondrial stress, and increased apoptotic priming. To determine whether these defects impacted NK cell persistence, we quantified the relative contribution of TiNK cells across defined windows of tumour microenvironment (TME) exposure.

The temporal structure of the TiNK compartment differed markedly between tumour types. In MC38 and E0771 tumours, 24 h, 48 h, and 72 h-labelled TiNK cells contributed equally to the overall population, consistent with sustained NK cell persistence. In contrast, B16 and LLC tumours were enriched for recently infiltrated cells, with progressive loss of 48 h and 72 h cohorts (Fig. 7E). These findings indicate reduced persistence of NK cells within B16 and LLC TMEs and a skewing toward newly infiltrating populations.

To determine whether this reflected altered recruitment or retention, we assessed expression of chemokine receptors and adhesion molecules. Key receptors implicated in NK cell trafficking, including Ccr2, Cxcr4, and Ccr5, as well as adhesion molecules such as LFA-1 and VLA-4, were maintained or increased in B16 and LLC TiNK cells. Consistent with this, overall NK cell density was comparable across tumour types (Fig. 7F-H), indicating that reduced persistence is unlikely to result from impaired recruitment.

We next tested whether enhancing interferon signalling could improve NK cell accumulation in tumours with low persistence. Adoptive transfer of IL-15-expanded NK cells^23,24^ stimulated with IL-18 plus IFN-β resulted in increased TiNK accumulation within B16 tumours compared with IL-18 stimulation alone, although tumour control was not improved (Fig. 7I-L).

Together, these data demonstrate that tumour-specific metabolic and translational stress programmes limit NK cell persistence within the TME. While MC38 and E0771 tumours support sustained NK cell residency, B16 and LLC tumours drive rapid loss of infiltrating NK cells, resulting in a temporal decay of the TiNK compartment. These findings identify impaired persistence, rather than reduced recruitment, as a defining feature of metabolically hostile tumour microenvironments, and suggest that strategies aimed at preserving NK cell biosynthetic capacity and stress resilience during early tumour exposure may enhance the durability of NK-based immunotherapies.

## Discussion

Using a controlled, multi-tumour comparative framework, this study demonstrates that tumour-infiltrating NK-cell dysfunction arises not from impaired recruitment or global nutrient limitation, but from an early failure of translational and mitochondrial homeostasis following tumour entry. These findings identify NK-cell persistence, rather than infiltration, as a key determinant of reduced NK cell abundance in B16 and LLC tumours and highlight translational regulation as a central constraint on NK-cell immunity within suppressive tumour microenvironments. While prior studies have examined how acute TME exposure (24-48 h) affect NK cell function at the transcriptomic or phenotypic level, they lacked direct resolution of NK-cell metabolic state and comparisons of distinct tumour TMEs^4,25^. By integrating proteomic, metabolomic, and time-resolved *in vivo* analyses across multiple tumour models, this study provides a systems-level view of NK cell dysfunction that is not accessible from single-tumour or transcriptomic approaches alone.

Despite implantation in identical anatomical sites, the four tumours examined imposed strikingly divergent NK-cell fates. MC38 and E0771 retained metabolically resilient and functionally competent NK populations, whereas B16 and LLC TMEs rapidly induced loss of effector function and mitochondrial integrity. Notably, within-tumour heterogeneity was most pronounced in B16, where translational repression, mitochondrial stress, and apoptotic priming varied markedly between samples, uggesting that heterogeneous suppressive niches within tumours may amplify NK dysfunction as tumours evolve.

Although NK-cell metabolism is often framed in terms of fuel availability and mTORC1 activation^3,8^, our proteomic analyses reveal that translational capacity itself represents a key limiting factor in suppressive TMEs^26,27^. B16 and LLC NK cells exhibited reduced abundance of key initiation factors and increased engagement of translational inhibitory mechanisms. Importantly, these defects emerged rapidly after tumour entry, indicating that translational repression is an early and active process rather than a downstream consequence of metabolic collapse.

NK cell dysfunction in tumours has often been described using concepts analogous to T cell exhaustion, characterised by progressive loss of effector function and altered receptor expression^28^. However, the application of “exhaustion” as a unifying framework for NK cells remains debated and may oversimplify the diversity of dysfunctional states that arise under chronic stimulation^29^. Our data suggest that, at least in these tumour models, NK cell impairment is more directly linked to disruption of core cellular homeostatic processes, particularly protein translation and mitochondrial maintenance^27^. This interpretation is consistent with growing evidence that metabolic and mitochondrial dysfunction are central drivers of NK cell failure within the tumour microenvironment^4,17^. Unlike classical exhaustion paradigms, which are frequently defined by transcriptional reprogramming and inhibitory receptor engagement, the dominant defects observed here emerge rapidly following tumour entry and are coupled to proteostatic stress and reduced biosynthetic capacity.

In addition, our data indicate selective disruption of the translation initiation machinery, most notably through reduced abundance of eIF1 in TiNK cells, particularly within B16 and LLC tumours. eIF1 plays a critical role in enforcing start codon selection fidelity during translation initiation, and reduced eIF1 levels have been shown to relax initiation stringency, promoting the use of non-canonical start sites and alternative upstream open reading frames^30,31^. This can result in substantial reprogramming of the cellular translatome, including proteins involved in energy metabolism and mitochondrial function, without requiring changes in mRNA abundance^32^. Notably, prior work has linked eIF1-dependent translational control to perturbations in cellular ATP levels and mitochondrial respiration, suggesting that altered initiation fidelity may directly influence metabolic capacity^32^. Together, these findings raise the possibility that reduced eIF1 in TiNK cells contributes not only to global translational suppression but also to metabolic dysfunction through selective reprogramming of the translated proteome. Importantly, this mode of regulation is mechanistically distinct from the integrated stress response, which primarily modulates global translation rates via eIF2 signalling. Together, these findings suggest that, in addition to translational suppression, TiNK cells may undergo qualitative reprogramming of protein synthesis, altering both the abundance and fidelity of translated proteins and thereby reshaping the functional proteome available to support effector function, stress adaptation, and survival.

Metabolomic profiling demonstrated that tumour microenvironments occupy distinct, structured metabolic states in the PCA space. However, these differences did not converge on a single metabolic feature explaining NK dysfunction. Instead, metabolic variation appears to provide a contextual backdrop that may influence cellular stress responses rather than directly determining NK cell fate.

Mitochondrial dysfunction in B16 and LLC NK cells was characterised by early loss of mitochondrial mass, selective engagement of mitochondrial stress-support pathways, and heightened dependence on mitochondrial and glycolytic flux. These features are consistent with a compensatory response to cellular stress^33,34^. The divergence between preserved membrane potential and reduced mass further supports a model in which mitochondrial stress is tolerated transiently but ultimately unsustainable in the context of chronic translational repression. Previous studies have described impairment of mitochondrial network in TiNK^16,35^. However, these measurements do not capture the dynamic metabolic adaptation of infiltrating NK cells, which in adverse TMEs, such as B16 and LLC, appears incomplete or unsuccessful.

Together, these findings support a model in which early translational repression acts upstream of broader functional decline. Impaired protein synthesis is likely to constrain the production of mitochondrial and cytoprotective proteins, thereby limiting the ability of NK cells to maintain mitochondrial integrity under stress. This coupling of translational capacity to mitochondrial fitness provides a mechanistic framework linking early proteostatic disruption to the later emergence of apoptotic priming and reduced persistence within the tumour microenvironment.

An additional observation from this study is that IFN-I signalling is associated with IL18r1 abundance across tumour models. Previous research has shown that IL-18 contributes to metabolic regulation and can support NK cell survival^36-38^. Here, our data suggest that IFN-I signalling may sustain NK cell fitness indirectly by preserving IL-18 responsiveness, thereby constraining apoptotic sensitivity rather than acutely boosting effector output.

Consistent with this interpretation, adoptive transfer experiments demonstrated that ex vivo exposure to IFN-I enhanced NK cell accumulation within established B16 tumours without improving short-term tumour control. This dissociation between increased NK cell numbers and unchanged tumour growth suggests that IFN-I priming primarily supports NK cell persistence within the tumour microenvironment rather than directly augmenting effector function. These observations align with the observed association between IFN-I signalling, IL18r1 expression, and reduced apoptotic priming, and support a model in which IFN-I contributes to NK cell fitness by preserving survival and stress tolerance in hostile tumour environments.

Several limitations should be considered. First, although proteomic and functional analyses strongly implicate translational repression and mitochondrial stress in NK cell dysfunction, direct causal relationships between these processes remain to be formally established. Second, while the multi-tumour framework provides a powerful comparative approach, the findings are restricted to transplantable murine models and may not fully capture the complexity of human tumours. Finally, although IFN-I signalling and IL-18 responsiveness were associated with NK cell fitness, the precise molecular mechanisms linking these pathways to translational control require further investigation.

These findings suggest that therapeutic strategies aimed at preserving translational capacity, proteostasis, and mitochondrial integrity, for example through modulation of stress response pathways or cytokine priming, may be more effective than approaches that solely enhance activation signals^39^. These findings suggest that strategies aimed at preserving NK cell biosynthetic capacity and stress resilience during early tumour exposure may enhance the durability of NK-based immunotherapies. Interventions applied early during tumour entry may be particularly important to prevent the establishment of irreversible dysfunction.

## Supplementary figure legends

**Supplementary Figure 1:**
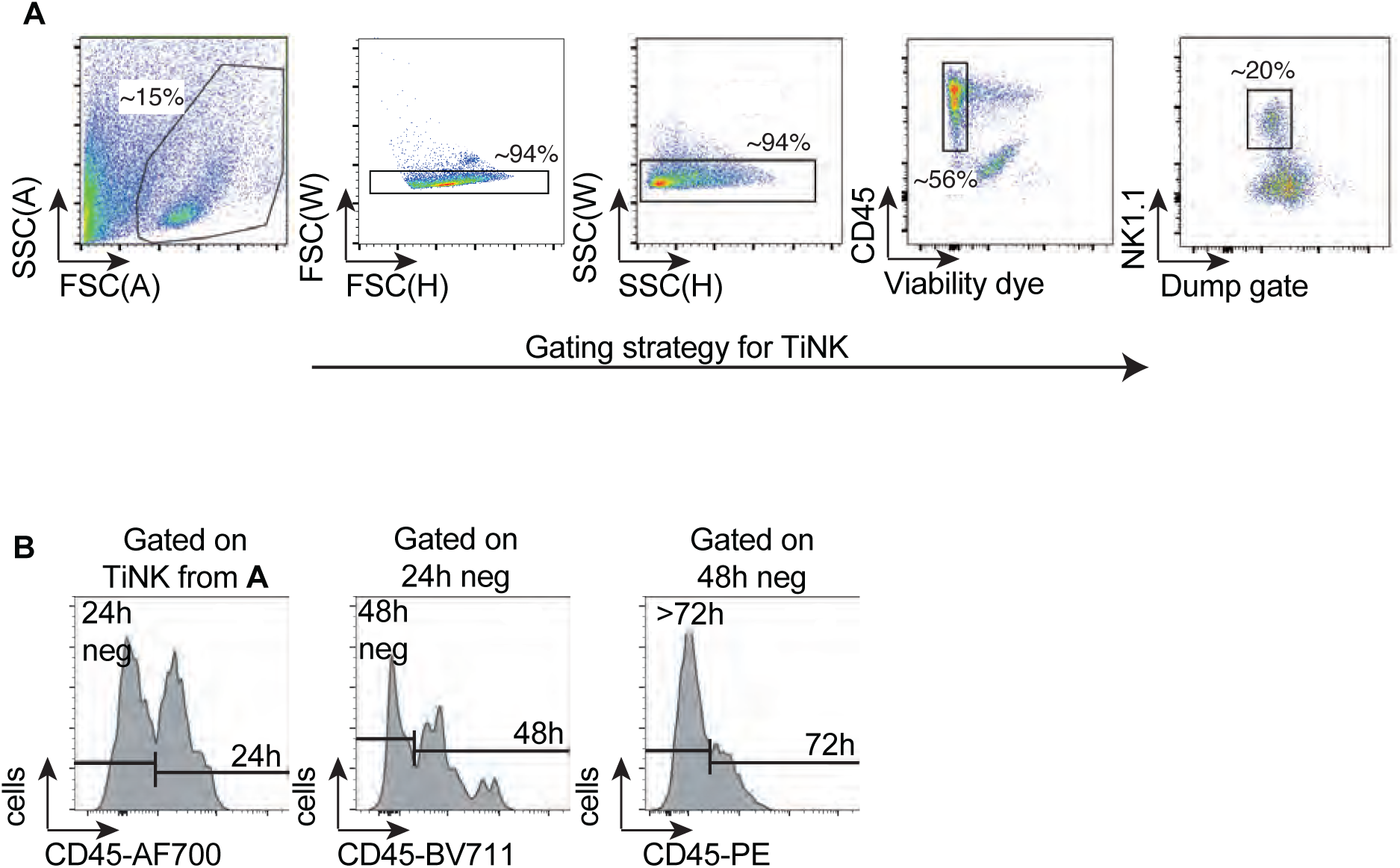
**(A,B)** Gating strategies for TiNK from processed tumours (A) and subsequent gating for time in tumour groups following time-stamping protocol (B).

**Supplementary Figure 2:**
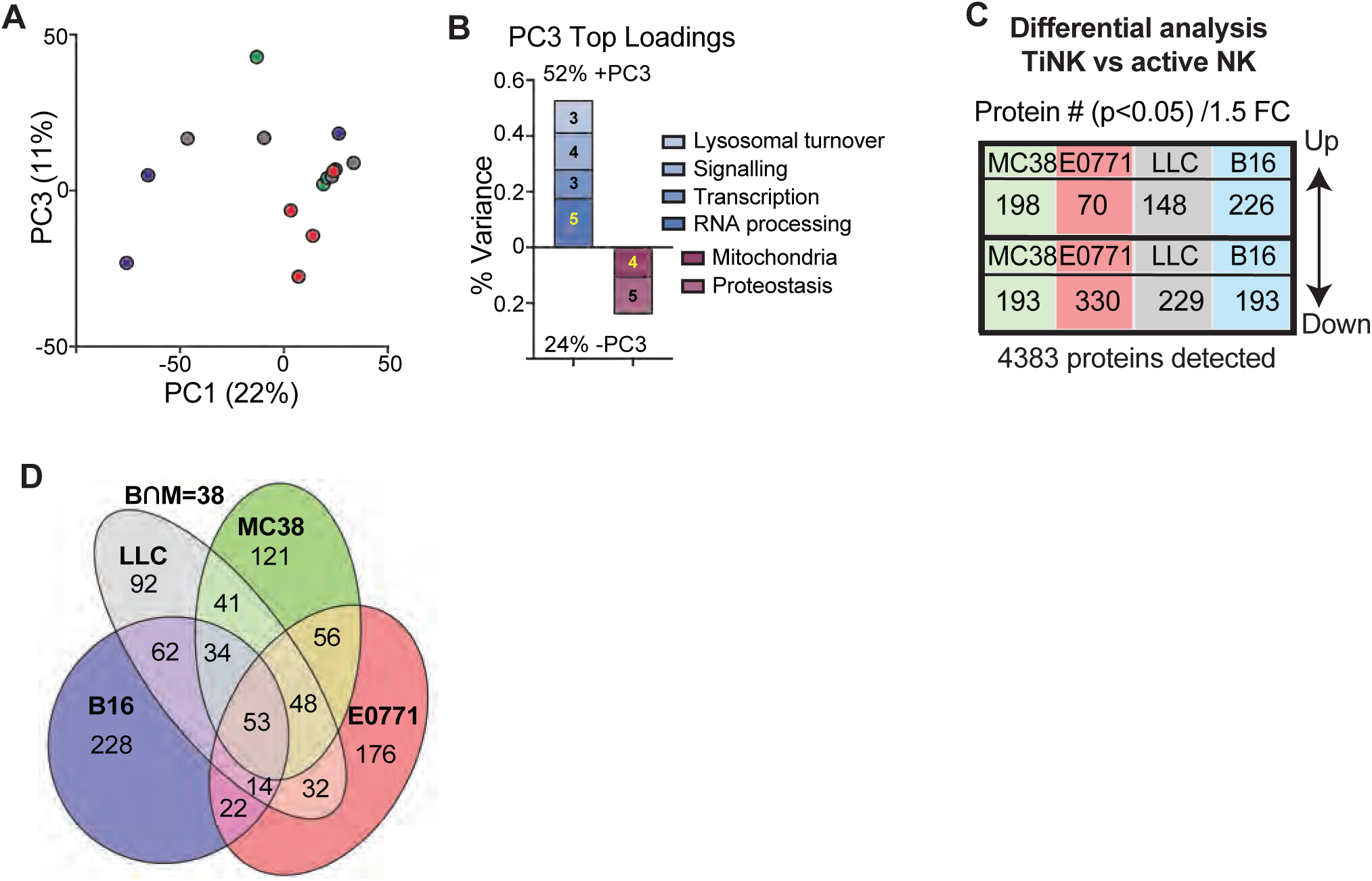
Mice were challenged with MC38 (green), B16 (blue), LLC (grey), or E0771 (red) tumours (s.c. 14-15 days). Mice injected with P Poly(I:C) for 24 hours were used to sort naïve and *in vivo* activated NK cells. Splenic NK and TiNK proteomes were quantified. (**A,B**) PCA analysis of TiNK cells showing PC1 versus PC3 (A) and the top 50 positive and negative loading proteins for PC3 (B). (**C**) Table summarising the DE proteins from the 4 parallel Limma analysis (Limma linear modelling, p<0.05, 1.5 fold change) of TiNK versus in vivo activated NK cells. (**D**) Venn diagram showing the overlapping proteins that we DE in each TiNK compared to in vivo activated NK cells. Overlap between B16 and MC38 is given in top right outside the ellipses. Data represent individual proteomic samples (n = 4 [per group) collected from a least 2 independent experiments.

**Supplementary Figure 3:**
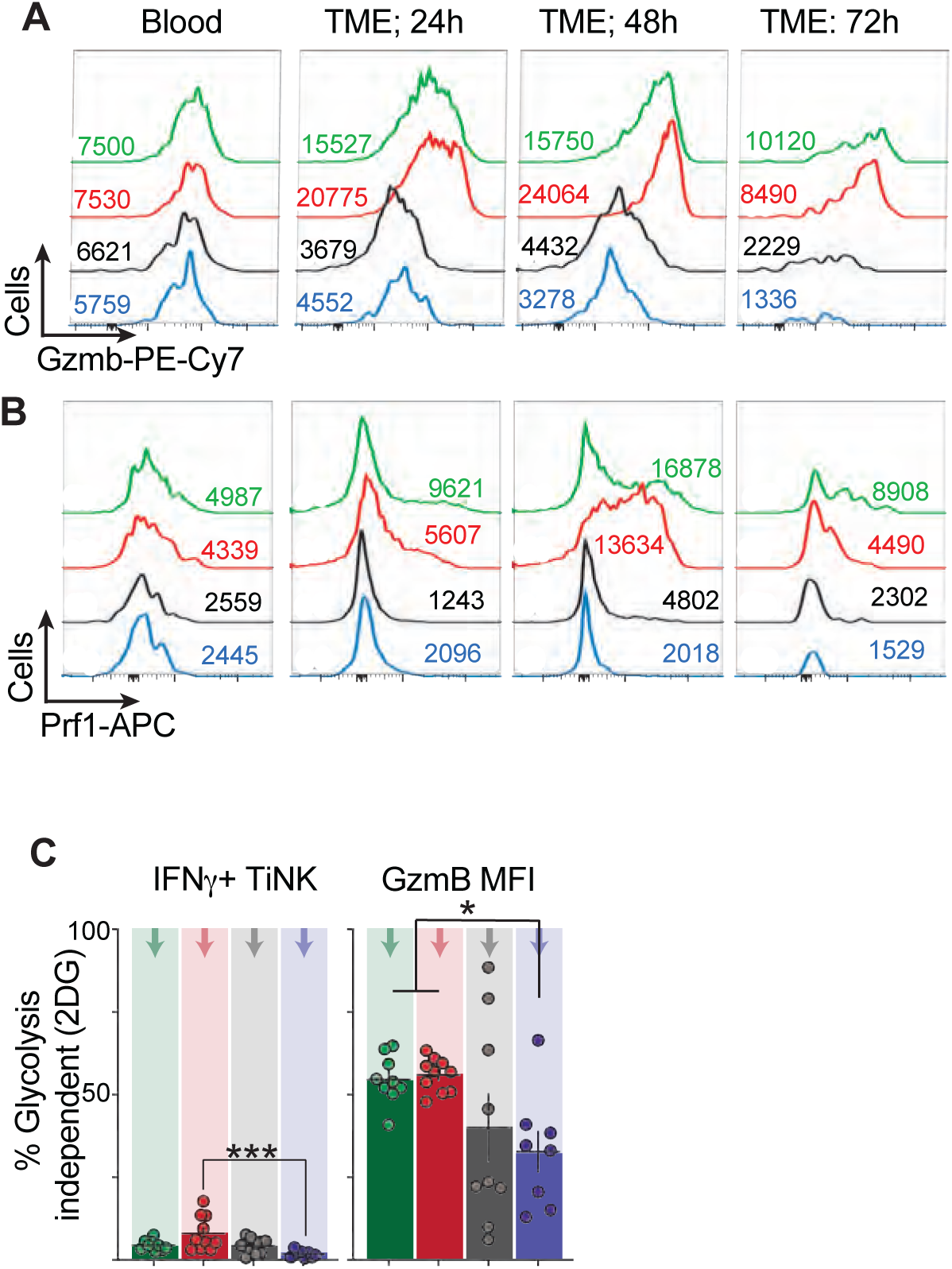
**(A-B)** Mice were challenged indicated tumours and analysed following time-stamping to quantify duration of TME exposure before harvesting tumours on day 15 and analysis by flow cytometry. Representative histograms with MFI values of Granzyme B (A) and Perforin (B) expression in the blood and after exposure to respective TME for 24-72 hours. **(C)** TiNK were isolated from day 14-15 tumours, stimulated for 3 h with PMA/ionomycin +/- 2-deoxyglucose (2DG, 50 mM)) and glycolysis independent IFNγ and granzyme B shown. Data represent individual mice (n = 8-9 [A,B] 8-10 [C] per group) from a least 2 independent experiments. Statistical analysis was performed using ANOVA or non-parametric tests as appropriate, with Tukey post-tests. ns, not significant; *p < 0.05; ***p < 0.001.

## Acknowledgements

MN and CFA were supported by funding from the Enterprise Ireland innovative partnership grant with ONK Therapeutics (IP20210976) and two Marie Skłodowska-Curie Actions fellowships (101111015 and 101108116). CC was supported by a Government of Ireland studentship (GOIPG/2022/81). DKF is supported by Research Ireland (IRCLA/2023/1402 and 22/FFP-A/10326). We thank Barry Moran and the flow cytometry facility for the technical support they provided during this research project.

## Declaration of generative AI and AI-assisted technologies in the manuscript preparation process

During the preparation of this manuscript, the authors used Copilot to assist with editing for clarity, grammar, and stylistic consistency, and to support troubleshooting of R code used for data analysis. The authors critically reviewed, edited, and validated all outputs generated by this tool and take full responsibility for the accuracy, integrity, and originality of the content of the published article.

## Data availability

The mass spectrometry proteomics data have been deposited to the [PRIDE/ProteomeXchange Consortium] with the dataset identifier [PXDXXXXXX]. The metabolomics raw data will be deposited in MetaboLights. The data will be released upon acceptance of the manuscript, and corresponding accession numbers will be included in the published article.

## Supplementary tables

**Supplementary Table 1:**
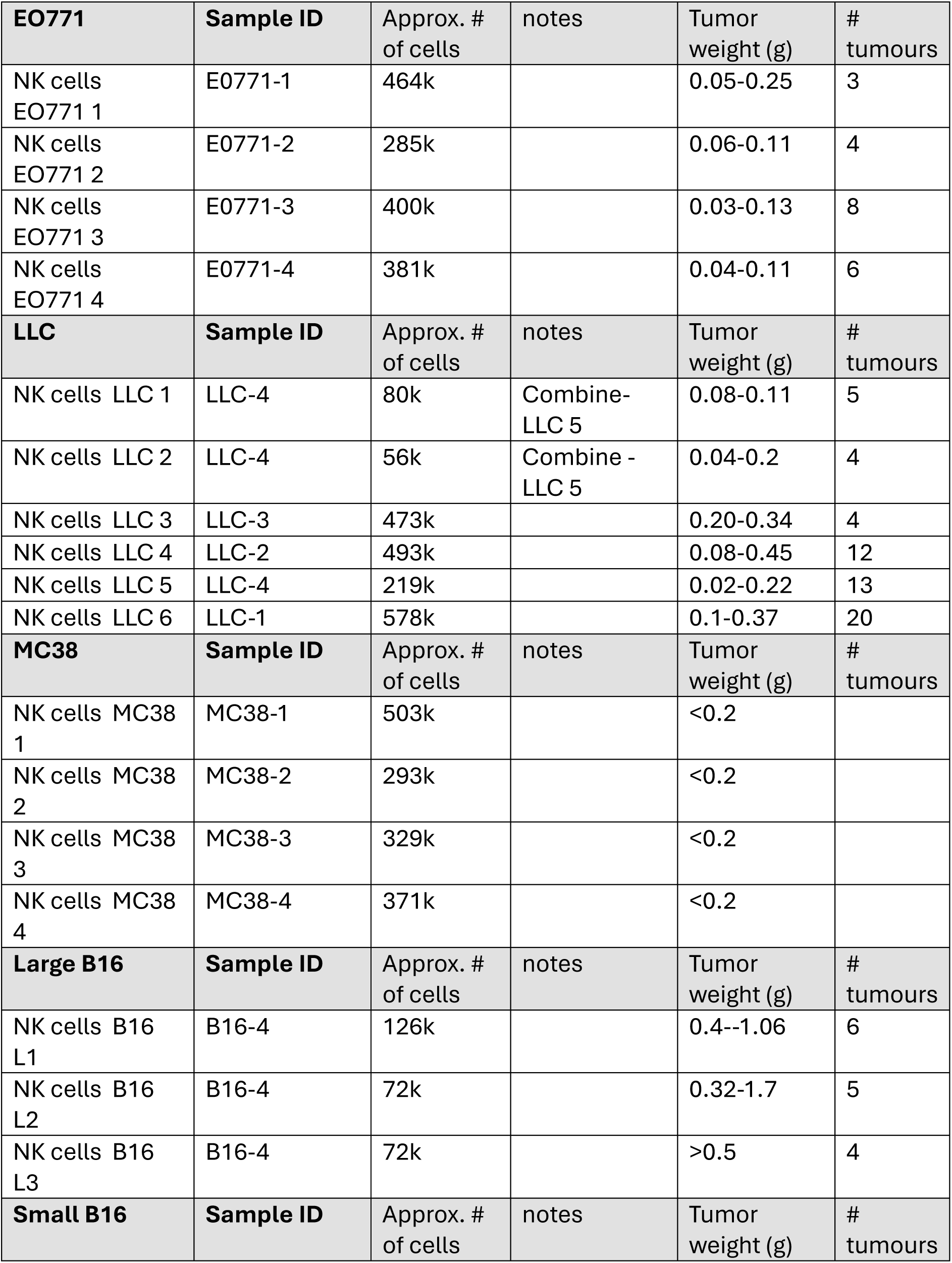
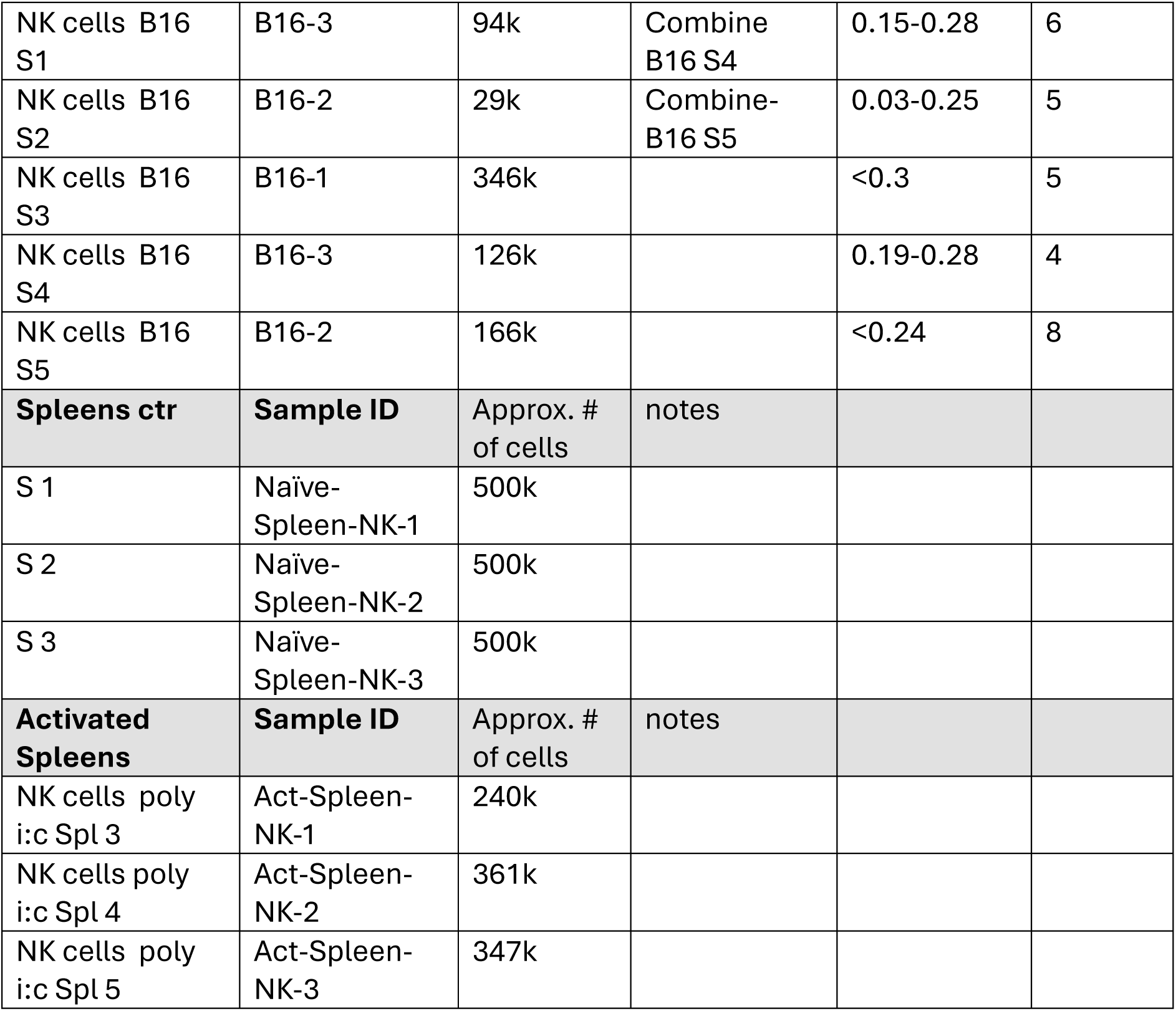
Proteomic samples.

| **EO771** | **Sample ID** | Approx. #<br>of cells | notes | Tumor<br>weight (g) | #<br>tumours |
| --- | --- | --- | --- | --- | --- |
| NK cells<br>EO771 1 | E0771-1 | 464k |  | 0.05-0.25 | 3 |
| NK cells<br>EO771 2 | E0771-2 | 285k |  | 0.06-0.11 | 4 |
| NK cells<br>EO771 3 | E0771-3 | 400k |  | 0.03-0.13 | 8 |
| NK cells<br>EO771 4 | E0771-4 | 381k |  | 0.04-0.11 | 6 |
| **LLC** | **Sample ID** | Approx. #<br>of cells | notes | Tumor<br>weight (g) | #<br>tumours |
| NK cells LLC 1 | LLC-4 | 80k | Combine-LLC 5 | 0.08-0.11 | 5 |
| NK cells LLC 2 | LLC-4 | 56k | Combine -<br>LLC 5 | 0.04-0.2 | 4 |
| NK cells LLC 3 | LLC-3 | 473k |  | 0.20-0.34 | 4 |
| NK cells LLC 4 | LLC-2 | 493k |  | 0.08-0.45 | 12 |
| NK cells LLC 5 | LLC-4 | 219k |  | 0.02-0.22 | 13 |
| NK cells LLC 6 | LLC-1 | 578k |  | 0.1-0.37 | 20 |
| **MC38** | **Sample ID** | Approx. #<br>of cells | notes | Tumor<br>weight (g) | #<br>tumours |
| NK cells MC38<br>1 | MC38-1 | 503k |  | <0.2 |  |
| NK cells MC38<br>2 | MC38-2 | 293k |  | <0.2 |  |
| NK cells MC38<br>3 | MC38-3 | 329k |  | <0.2 |  |
| NK cells MC38<br>4 | MC38-4 | 371k |  | <0.2 |  |
| **Large B16** | **Sample ID** | Approx. #<br>of cells | notes | Tumor<br>weight (g) | #<br>tumours |
| NK cells B16<br>L1 | B16-4 | 126k |  | 0.4--1.06 | 6 |
| NK cells B16<br>L2 | B16-4 | 72k |  | 0.32-1.7 | 5 |
| NK cells B16<br>L3 | B16-4 | 72k |  | >0.5 | 4 |
| **Small B16** | **Sample ID** | Approx. #<br>of cells | notes | Tumor<br>weight (g) | #<br>tumours |
| NK cells B16 S1 | B16-3 | 94k | Combine B16 S4 | 0.15-0.28 | 6 |
| NK cells B16 S2 | B16-2 | 29k | Combine-B16 S5 | 0.03-0.25 | 5 |
| NK cells B16 S3 | B16-1 | 346k |  | <0.3 | 5 |
| NK cells B16 S4 | B16-3 | 126k |  | 0.19-0.28 | 4 |
| NK cells B16 S5 | B16-2 | 166k |  | <0.24 | 8 |
| **Spleens ctr** | **Sample ID** | Approx. # of cells | notes |  |  |
| S 1 | Naïve-Spleen-NK-1 | 500k |  |  |  |
| S 2 | Naïve-Spleen-NK-2 | 500k |  |  |  |
| S 3 | Naïve-Spleen-NK-3 | 500k |  |  |  |
| **Activated Spleens** | **Sample ID** | Approx. # of cells | notes |  |  |
| NK cells poly i:c Spl 3 | Act-Spleen-NK-1 | 240k |  |  |  |
| NK cells poly i:c Spl 4 | Act-Spleen-NK-2 | 361k |  |  |  |
| NK cells poly i:c Spl 5 | Act-Spleen-NK-3 | 347k |  |  |  |

**Supplementary table 2:** list of antibodies.

| Target | Clone | Fluorophore(s) | Vendor |
| --- | --- | --- | --- |
| CD3 | 17A2 | BV510, BV650 | BioLegend |
| CD19 | 6D5, ID3 | BV510, BV650, APC | BioLegend, BD |
| Ly6G | 1A8 | BV510 | BioLegend |
| F4/80 | BM8 | BV510, PECy7 | BioLegend |
| CD11b | M1/70 | BV605 | BioLegend |
| NK1.1 | PK136 | PE-Cy5, BV785 | BioLegend |
| CD45 | 30-F11 | For analysis: PE, FITC<br>For time stamping: PE, R718, BV711 | BioLegend, BD, ThermoFisher |
| CD69 | H1.2F3 | BV650, PE-Dazzle594 | BioLegend |
| IFNγ | XMG1.2 | BV421 | BioLegend |
| TNF | MP6-XT22 | BV711 | BD |
| Granzyme B | NGZB | PE-Cy7 | ThermoFisher |
| Perforin | S16009B | APC | BioLegend |

**Supplementary table 3:**
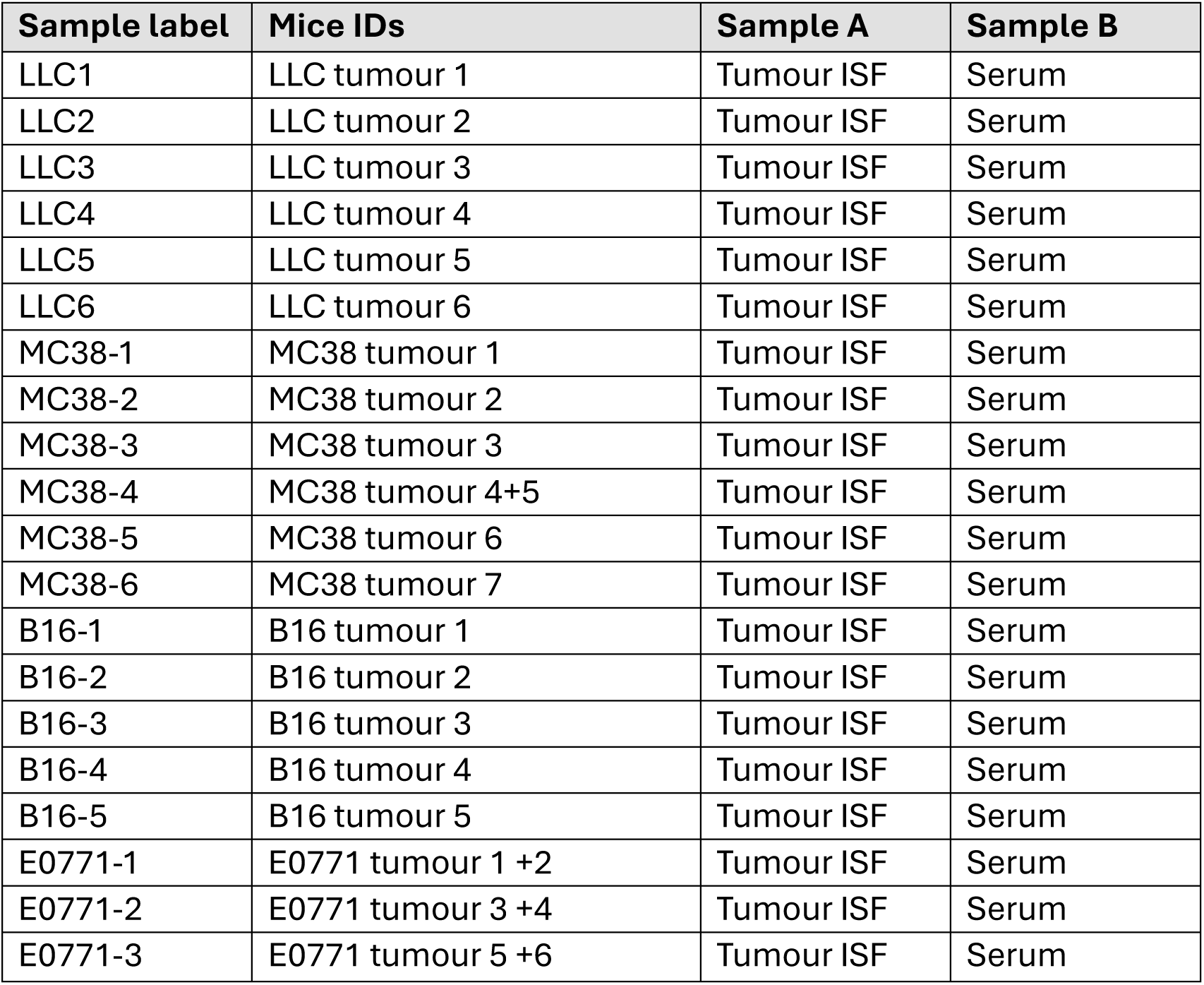
Metabolomics samples.

| Sample label | Mice IDs | Sample A | Sample B |
| --- | --- | --- | --- |
| LLC1 | LLC tumour 1 | Tumour ISF | Serum |
| LLC2 | LLC tumour 2 | Tumour ISF | Serum |
| LLC3 | LLC tumour 3 | Tumour ISF | Serum |
| LLC4 | LLC tumour 4 | Tumour ISF | Serum |
| LLC5 | LLC tumour 5 | Tumour ISF | Serum |
| LLC6 | LLC tumour 6 | Tumour ISF | Serum |
| MC38-1 | MC38 tumour 1 | Tumour ISF | Serum |
| MC38-2 | MC38 tumour 2 | Tumour ISF | Serum |
| MC38-3 | MC38 tumour 3 | Tumour ISF | Serum |
| MC38-4 | MC38 tumour 4+5 | Tumour ISF | Serum |
| MC38-5 | MC38 tumour 6 | Tumour ISF | Serum |
| MC38-6 | MC38 tumour 7 | Tumour ISF | Serum |
| B16-1 | B16 tumour 1 | Tumour ISF | Serum |
| B16-2 | B16 tumour 2 | Tumour ISF | Serum |
| B16-3 | B16 tumour 3 | Tumour ISF | Serum |
| B16-4 | B16 tumour 4 | Tumour ISF | Serum |
| B16-5 | B16 tumour 5 | Tumour ISF | Serum |
| E0771-1 | E0771 tumour 1 +2 | Tumour ISF | Serum |
| E0771-2 | E0771 tumour 3 +4 | Tumour ISF | Serum |
| E0771-3 | E0771 tumour 5 +6 | Tumour ISF | Serum |

## References

1. Vivier, E., Tomasello, E., Baratin, M., Walzer, T., and Ugolini, S. (2008). Functions of natural killer cells. Nat Immunol 9, 503–510. 10.1038/ni1582.

2. Long, E.O., Kim, H.S., Liu, D., Peterson, M.E., and Rajagopalan, S. (2013). Controlling natural killer cell responses: integration of signals for activation and inhibition. Annu Rev Immunol 31, 227–258. 10.1146/annurev-immunol-020711-075005.

3. Kedia-Mehta, N., and Finlay, D.K. (2019). Competition for nutrients and its role in controlling immune responses. Nature communications 10, 2123. 10.1038/s41467-019-10015-4.

4. Dean, I., Lee, C.Y.C., Tuong, Z.K., Li, Z., Tibbitt, C.A., Willis, C., Gaspal, F., Kennedy, B.C., Matei-Rascu, V., Fiancette, R., et al. (2024). Rapid functional impairment of natural killer cells following tumor entry limits anti-tumor immunity. Nature communications 15, 683. 10.1038/s41467-024-44789-z.

5. Guillerey, C. (2020). NK Cells in the Tumor Microenvironment. Adv Exp Med Biol 1273, 69–90. 10.1007/978-3-030-49270-0_4.

6. Kennedy, P.R., Arvindam, U.S., Phung, S.K., Ettestad, B., Feng, X., Li, Y., Kile, Q.M., Hinderlie, P., Khaw, M., Huang, R.S., et al. (2024). Metabolic programs drive function of therapeutic NK cells in hypoxic tumor environments. Sci Adv 10, eadn1849. 10.1126/sciadv.adn1849.

7. Binnewies, M., Roberts, E.W., Kersten, K., Chan, V., Fearon, D.F., Merad, M., Coussens, L.M., Gabrilovich, D.I., Ostrand-Rosenberg, S., Hedrick, C.C., et al. (2018). Understanding the tumor immune microenvironment (TIME) for effective therapy. Nat Med 24, 541–550. 10.1038/s41591-018-0014-x.

8. Loftus, R.M., Assmann, N., Kedia-Mehta, N., O’Brien, K.L., Garcia, A., Gillespie, C., Hukelmann, J.L., Oefner, P.J., Lamond, A.I., Gardiner, C.M., et al. (2018). Amino acid-dependent cMyc expression is essential for NK cell metabolic and functional responses in mice. Nature communications 9, 2341. 10.1038/s41467-018-04719-2.

9. Donnelly, R.P., Loftus, R.M., Keating, S.E., Liou, K.T., Biron, C.A., Gardiner, C.M., and Finlay, D.K. (2014). mTORC1-dependent metabolic reprogramming is a prerequisite for NK cell effector function. J Immunol 193, 4477–4484. 10.4049/jimmunol.1401558.

10. Slattery, K., Woods, E., Zaiatz-Bittencourt, V., Marks, S., Chew, S., Conroy, M., Goggin, C., MacEochagain, C., Kennedy, J., Lucas, S., et al. (2021). TGFbeta drives NK cell metabolic dysfunction in human metastatic breast cancer. J Immunother Cancer 9. 10.1136/jitc-2020-002044.

11. Viel, S., Marcais, A., Guimaraes, F.S., Loftus, R., Rabilloud, J., Grau, M., Degouve, S., Djebali, S., Sanlaville, A., Charrier, E., et al. (2016). TGF-beta inhibits the activation and functions of NK cells by repressing the mTOR pathway. Sci Signal 9, ra19. 10.1126/scisignal.aad1884.

12. Rodriguez, P.C., Quiceno, D.G., Zabaleta, J., Ortiz, B., Zea, A.H., Piazuelo, M.B., Delgado, A., Correa, P., Brayer, J., Sotomayor, E.M., et al. (2004). Arginase I production in the tumor microenvironment by mature myeloid cells inhibits T-cell receptor expression and antigen-specific T-cell responses. Cancer research 64, 5839–5849. 10.1158/0008-5472.CAN-04-0465.

13. Novick, D. (2024). IL-18 and IL-18BP: A Unique Dyad in Health and Disease. Int J Mol Sci 25. 10.3390/ijms252413505.

14. Chambers, A.M., Wang, J., Lupo, K.B., Yu, H., Atallah Lanman, N.M., and Matosevic, S. (2018). Adenosinergic Signaling Alters Natural Killer Cell Functional Responses. Front Immunol 9, 2533. 10.3389/fimmu.2018.02533.

15. Ji, J.H., Armstrong, W.R., Li, J.H., Lalani, A., Lee, C.D., Bharadwaj, V., Ho, K., Liang, J., Muir, A., and O’Sullivan, T.E. (2026). The transcriptional repressor Fli1 inhibits proteostasis during nutrient stress to limit NK cell persistence in solid tumors. Immunity 59, 717–733 e710. 10.1016/j.immuni.2026.01.017.

16. Zheng, X., Qian, Y., Fu, B., Jiao, D., Jiang, Y., Chen, P., Shen, Y., Zhang, H., Sun, R., Tian, Z., and Wei, H. (2019). Mitochondrial fragmentation limits NK cell-based tumor immunosurveillance. Nat Immunol 20, 1656–1667. 10.1038/s41590-019-0511-1.

17. Poznanski, S.M., Singh, K., Ritchie, T.M., Aguiar, J.A., Fan, I.Y., Portillo, A.L., Rojas, E.A., Vahedi, F., El-Sayes, A., Xing, S., et al. (2021). Metabolic flexibility determines human NK cell functional fate in the tumor microenvironment. Cell Metab 33, 1205–1220 e1205. 10.1016/j.cmet.2021.03.023.

18. Pelgrom, L.R., Davis, G.M., O’Shaughnessy, S., Wezenberg, E.J.M., Van Kasteren, S.I., Finlay, D.K., and Sinclair, L.V. (2023). QUAS-R: An SLC1A5-mediated glutamine uptake assay with single-cell resolution reveals metabolic heterogeneity with immune populations. Cell Rep 42, 112828. 10.1016/j.celrep.2023.112828.

19. Akazawa, T., Ebihara, T., Okuno, M., Okuda, Y., Shingai, M., Tsujimura, K., Takahashi, T., Ikawa, M., Okabe, M., Inoue, N., et al. (2007). Antitumor NK activation induced by the Toll-like receptor 3-TICAM-1 (TRIF) pathway in myeloid dendritic cells. Proc Natl Acad Sci U S A 104, 252–257. 10.1073/pnas.0605978104.

20. Sareneva, T., Julkunen, I., and Matikainen, S. (2000). IFN-alpha and IL-12 induce IL-18 receptor gene expression in human NK and T cells. J Immunol 165, 1933–1938. 10.4049/jimmunol.165.4.1933.

21. Lee, A.J., Chen, B., Chew, M.V., Barra, N.G., Shenouda, M.M., Nham, T., van Rooijen, N., Jordana, M., Mossman, K.L., Schreiber, R.D., et al. (2017). Inflammatory monocytes require type I interferon receptor signaling to activate NK cells via IL-18 during a mucosal viral infection. J Exp Med 214, 1153–1167. 10.1084/jem.20160880.

22. Mattei, F., Schiavoni, G., Belardelli, F., and Tough, D.F. (2001). IL-15 is expressed by dendritic cells in response to type I IFN, double-stranded RNA, or lipopolysaccharide and promotes dendritic cell activation. J Immunol 167, 1179–1187. 10.4049/jimmunol.167.3.1179.

23. Assmann, N., O’Brien, K.L., Donnelly, R.P., Dyck, L., Zaiatz-Bittencourt, V., Loftus, R.M., Heinrich, P., Oefner, P.J., Lynch, L., Gardiner, C.M., et al. (2017). Srebp-controlled glucose metabolism is essential for NK cell functional responses. Nat Immunol 18, 1197–1206. 10.1038/ni.3838.

24. Michelet, X., Dyck, L., Hogan, A., Loftus, R.M., Duquette, D., Wei, K., Beyaz, S., Tavakkoli, A., Foley, C., Donnelly, R., et al. (2018). Metabolic reprogramming of natural killer cells in obesity limits antitumor responses. Nat Immunol 19, 1330–1340. 10.1038/s41590-018-0251-7.

25. Kirschenbaum, D., Xie, K., Ingelfinger, F., Katzenelenbogen, Y., Abadie, K., Look, T., Sheban, F., Phan, T.S., Li, B., Zwicky, P., et al. (2024). Time-resolved single-cell transcriptomics defines immune trajectories in glioblastoma. Cell 187, 149–165 e123. 10.1016/j.cell.2023.11.032.

26. Suresh, S., and O’Donnell, K.A. (2021). Translational Control of Immune Evasion in Cancer. Trends Cancer 7, 580–582. 10.1016/j.trecan.2021.04.002.

27. Piccirillo, C.A., Bjur, E., Topisirovic, I., Sonenberg, N., and Larsson, O. (2014). Translational control of immune responses: from transcripts to translatomes. Nat Immunol 15, 503–511. 10.1038/ni.2891.

28. Bi, J., and Tian, Z. (2017). NK Cell Exhaustion. Front Immunol 8, 760. 10.3389/fimmu.2017.00760.

29. Murphy, W.J. (2025). Exhaustion is getting complicated. Blood 146, 3008–3010. 10.1182/blood.2025031002.

30. Ivanov, I.P., Loughran, G., Sachs, M.S., and Atkins, J.F. (2010). Initiation context modulates autoregulation of eukaryotic translation initiation factor 1 (eIF1). Proceedings of the National Academy of Sciences of the United States of America 107, 18056–18060. 10.1073/pnas.1009269107.

31. Grosely, R., Alvarado, C., Ivanov, I.P., Nicholson, O.B., Puglisi, J.D., Dever, T.E., and Lapointe, C.P. (2025). eIF1 and eIF5 dynamically control translation start site fidelity. Nat Struct Mol Biol 32, 2308–2318. 10.1038/s41594-025-01629-y.

32. Fijalkowska, D., Verbruggen, S., Ndah, E., Jonckheere, V., Menschaert, G., and Van Damme, P. (2017). eIF1 modulates the recognition of suboptimal translation initiation sites and steers gene expression via uORFs. Nucleic acids research 45, 7997–8013. 10.1093/nar/gkx469.

33. Cheng, Y.W., Liu, J., and Finkel, T. (2023). Mitohormesis. Cell Metab 35, 1872–1886. 10.1016/j.cmet.2023.10.011.

34. Kim, S., Ramalho, T.R., and Haynes, C.M. (2024). Regulation of proteostasis and innate immunity via mitochondria-nuclear communication. J Cell Biol 223. 10.1083/jcb.202310005.

35. Tumino, N., Nava Lauson, C.B., Tiberti, S., Besi, F., Martini, S., Fiore, P.F., Scodamaglia, F., Mingari, M.C., Moretta, L., Manzo, T., and Vacca, P. (2023). The tumor microenvironment drives NK cell metabolic dysfunction leading to impaired antitumor activity. Int J Cancer 152, 1698–1706. 10.1002/ijc.34389.

36. Ni, F., Zhang, T., Xiao, W., Dong, H., Gao, J., Liu, Y., and Li, J. (2021). IL-18-Mediated SLC7A5 Overexpression Enhances Osteogenic Differentiation of Human Bone Marrow Mesenchymal Stem Cells via the c-MYC Pathway. Front Cell Dev Biol 9, 748831. 10.3389/fcell.2021.748831.

37. Li, W., Jin, D., Takai, S., Inoue, N., Yamanishi, K., Tanaka, Y., and Okamura, H. (2024). IL-18 primes T cells with an antigen-inexperienced memory phenotype for proliferation and differentiation into effector cells through Notch signaling. Journal of leukocyte biology 117. 10.1093/jleuko/qiae172.

38. Hodge, D.L., Subleski, J.J., Reynolds, D.A., Buschman, M.D., Schill, W.B., Burkett, M.W., Malyguine, A.M., and Young, H.A. (2006). The proinflammatory cytokine interleukin-18 alters multiple signaling pathways to inhibit natural killer cell death. J Interferon Cytokine Res 26, 706–718. 10.1089/jir.2006.26.706.

39. Page, A., Chuvin, N., Valladeau-Guilemond, J., and Depil, S. (2024). Development of NK cell-based cancer immunotherapies through receptor engineering. Cell Mol Immunol 21, 315–331. 10.1038/s41423-024-01145-x.

